# Composition variations in archaeological human bone proteomes

**DOI:** 10.1101/2025.06.02.657369

**Authors:** Ragnheiður Diljá Ásmundsdóttir, Gaudry Troché, Jesper V. Olsen, Sarah Schrader, Frido Welker

## Abstract

Sampling strategies within the field of skeletal palaeoproteomics are often based on specimen availability. Knowledge of bone biology might assist in improving sample selection strategies and minimise unnecessary sampling of precious (hominin) material. We study ten bone sample locations across four bone elements, for a total of 10 adult, archaeological human skeletons. We compare bone proteome composition and modification for skeletal elements formed through endochondral and intramembranous ossification, as well as cortical-trabecular bone pairs of three skeletal locations. We observe minimal differences in bones formed through the two ossification processes, outside of the exclusive presence of cartilage-related proteins in endochondral bone samples. We observe higher protein concentrations, a larger number of protein groups and peptides, and lower rates of deamidation in cortical bone compared to trabecular bone proteomes, this indicates that cortical bone provides a better preservation environment compared to trabecular bone. Throughout our analysis, the petrous bone stands out, with the largest and most complex proteomes recovered for all studied individuals. Formed through endochondral ossification, the petrous bone undergoes minimal turnover during life. Our observations indicate that the petrous bone is the ideal source of ancient protein sequence information.

## 1. Introduction

The human skeleton is largely composed of three distinct proteomes: dental enamel, dentine, and bone proteomes, with the bone proteome being the most extensive consisting of over a thousand proteins in the living bone (Alves et al., 2011). The bone proteome, alongside the dental ones, is frequently used in human and faunal evolutionary research as it contains a few proteins with known amino acid substitutions that are informative in phylogenetic contexts. Furthermore, archaeological bone proteomes have been analysed to investigate past health status through the analysis of host proteins (Sawafuji et al., 2017) as well as proteins deriving from infectious disease agents (Wilkin et al., 2024). With the limited nature of the hominin fossil record in mind, great care needs to be taken to minimise destructive and/or unnecessary sampling.

Palaeoproteomic analyses are inherently based on a destructive sampling, and care should therefore be taken to determine sampling locations for optimal proteome recovery. In addition, in order to obtain comparable data across specimen cohorts for health-related palaeoproteomic and medical studies, chosen sampling locations require comparability in terms of composition and preservation in healthy, unaffected individuals. In other ancient biomolecular fields, such as ancient DNA research and isotopic research, researchers have optimised their sampling selection strategies (Alberti et al., 2018; Damgaard et al., 2015; Hansen et al., 2017; Harvig et al., 2014; R. Hedges et al., 2008; Jørkov et al., 2009; Kontopoulos et al., 2019; Parker et al., 2020; Pinhasi et al., 2015). This has not been done yet within the field of palaeoproteomics, where sampling selection strategies are generally based on specimen availability. As a result, in palaeoproteomics, it is generally expected for the bone proteome to be uniform across the skeleton, implying that the same proteome will be acquired regardless of utilised sampling location. This is, however, unlikely to be the case based on the biology of the living bone.

Bones in the human body are formed through two ossification processes, endochondral and intramembranous ossification. The two processes take place in different areas of the body and form bone with different structures. Endochondral ossification is found in the majority of the skeleton, from the base of the skull down the axial skeleton, the appendicular skeleton, as well as the petrous pyramid of the temporal bone (Jørkov et al., 2009; E. J. Mackie et al., 2011). It starts with mesenchymal stem cells at the location of the forming bone that are differentiated into chondrocytes that form a cartilage scaffold of the forming bone. This scaffold is then gradually ossified and replaced by bone tissue, and in this process chondrocytes are replaced by osteoblasts (Galea et al., 2021; Hallett et al., 2021; E. J. Mackie et al., 2008). In contrast, intramembranous ossification takes place mainly in the craniofacial skeleton, forming the flat bones of the face and cranium, and starts with mesenchymal stem cells that are differentiated directly into osteoblasts. Here, the ossification occurs within the soft tissue surrounding the location of the forming bone, without a cartilage scaffold (Celik et al., 2021; Galea et al., 2021; E. J. Mackie et al., 2011).

The different cellular composition of the two ossification processes are likely to influence the bone proteome. In endochondral ossification there are cartilage-related proteins present, such as collagen type X (COL10A1) (Alini et al., 1996; Deng et al., 2018), aggrecan core protein (ACAN) (Wilson et al., 2008), and cartilage oligomeric matrix protein (COMP) (Hedbom et al., 1992; Roughley, 2001), during the initial formation and until the bone is fully ossified. These proteins should therefore be absent in the proteomes resulting from intramembranous ossification. A few proteins, such as collagen type 2 (COL2A1) and bone morphogenic proteins (BMPs), are found in both cartilage and bone tissues (Deng et al., 2018). The age at which initial bone ossification is completed varies across the skeleton and between individuals, with most bones fully mineralised between late adolescence (for example the distal femoral epiphysis (Daghighi et al., 2021; Galić et al., 2016)), and the mid-second decade of life (for example the clavicles (Ekizoglu et al., 2015; Schulz et al., 2008)).

The majority of bones within the skeleton are composed of two different types of bone, cortical and trabecular bone. These two bones differ in terms of structure and function, with cortical bone having higher mineral as well as lower water content compared to trabecular bone (Beresheim et al., 2020; Faraldi et al., 2022; Gong et al., 1964; Haverfield et al., 2023; Lehtinen et al., 2004). The two bone types also differ in terms of maintenance during life. Bone turnover, the process of replacing the mineralised tissue (Sansalone et al., 2021), occurs at a much faster rate in trabecular bone (20-30% per year) compared to cortical bone (3-10% per year) (Deftos, 1998; Parfitt, 1994, 2002; Parfitt et al., 1996). How bone turnover affects proteome composition, especially the variation related to the ossification processes with the presence of cartilage-related proteins, in the adult bone, modern or archaeological, is unclear.

To optimise palaeoproteomic bone sampling selection strategies it is important to gain an understanding of the influences of bone biology and protein preservation on archaeological bone proteome composition. To do so we focus on two aspects of bone biology of the living bone: initial bone formation and bone maintenance during life. To what extent the differences are preserved in archaeological bone remains unclear. The structural differences between cortical and trabecular bone have previously been shown to result in differences in proteome preservation, with trabecular bone proteomes being smaller and containing more advanced degradation compared to cortical bone (Ásmundsdóttir et al., 2024). How the biology of the living bone influences the bone proteome in human archaeological material has previously not been extensively studied.

Here we compare the human bone proteome in terms of proteome size, composition and degradation to gain insight into the effects of bone biology on the archaeological proteome. We observe minimal differences in proteome size in bones formed through the two ossification processes in adult individuals. The main differences between the ossification processes are observed in the proteome composition, where cartilage-related proteins are only observed in adult endochondral bone. The petrous pyramid contains the greatest proteome among all sample locations, due to its minimal bone turnover during life. Cortical bone proteomes are larger and less degraded than the trabecular bone proteomes, consistent with previous studies.

## 2. Materials and methods

### 2.1. Archaeological material

In this study four bones from ten individuals were analysed. The skeletons originate from the Saint Eusebius church graveyard, Arnhem, the Netherlands and are currently housed at Leiden University, the Netherlands. The skeletons belonged to four females and six males and were selected based on the following criteria: they belonged to individuals ranging in age from late- young adults to middle age, and who showed no bone pathologies. The biological sex and age were estimated using bone morphology using standard osteological methods (Buikstra & Ubelaker, 1994), prior to this study. The individuals date to the mediaeval (ca. 1350 - 1650) and post-mediaeval (ca. 1650 - 1829) periods (Table 1). The two burial periods are estimated based on material finds in the soil layer underneath the graves, dateable finds associated with the graves (such as coins), and radiocarbon dating of selected graves within the cemetery (n=19). The graves were located in a homogenised brown, humus-containing layer (Zielman & Baetsen, 2020).

**Table 1.** Archaeological samples in this study. Biological sex estimation is based on common osteoarchaeological assessment.

| ID | Est. biological sex | Est. age range (in years) | Burial period (years AD) |
| --- | --- | --- | --- |
| V0868 | Male | Middle adult (36-49) | 1350-1650 |
| V1428 | Male | Middle adult (36-49) | 1650-1829 |
| V1506 | Female | Middle adult (36-49) | 1350-1650 |
| V1621 | Male | Adult (18-35) | 1650-1829 |
| V2141 | Male | Adult (18-35) | 1350-1650 |
| V2190 | Male | Adult (18-35) | 1350-1650 |
| V2193 | Female | Adult (18-35) | 1350-1650 |
| V2324 | Female | Adult (18-35) | 1350-1650 |
| V2351 | Female | Middle adult (36-49) | 1350-1650 |
| V2406 | Male | Middle adult (36-49) | 1350-1650 |

From each individual a rib and a femur were chosen to represent bones formed through endochondral ossification, and a parietal bone to represent bones formed through intramembranous ossification. Cortical and trabecular tissues were sampled from each of these three bone elements. From the distal femur, an additional sample was taken to represent a growth line. From the parietal bone, an additional sample was taken from a suture, when present, as well as an extra cortical bone sample from the interior side (exterior side was sampled in the description above). Finally, the petrous pyramid of the temporal bone was sampled as well. The temporal bone is a complex bone in terms of ossification, where the squamous and tympanic portions are formed through intramembranous ossification while the petrous portion is formed through endochondral ossification (Tubbs et al., 2012). The petrous pyramid undergoes minimal skeletal remodelling after initial ossification (Harvig et al., 2014; Jørkov et al., 2009). It is also an exceptionally dense skeletal element (Kontopoulos et al., 2019) that has proven to be an excellent source of ancient DNA (Alberti et al., 2018; Damgaard et al., 2015; Hansen et al., 2017; Parker et al., 2020; Pinhasi et al., 2015).

### 2.2. Methods

#### 2.2.1. Protein extraction

Bone samples were taken from each of the ten skeletons, with the exception of cranial sutures which were absent in two parietal bone specimens. The bone samples were acquired using diamond drill bits in a handheld drill. The drill and drill bit, as well as the sampling hood, were cleaned with 5% bleach solution and 70% ethanol between every sample. For each location 20.54 ± 1.11 mg bone powder (see exact amount per sample in SI Table 1) were placed in microtubes (Protein LoBind Tubes, Eppendorf) for extraction. A laboratory blank was included for each skeleton.

Protein extraction was performed using a modified protocol described in (Lanigan et al., 2020). Bone powder was demineralised using 1 mL UltraPure 0.5 M EDTA pH 8.0 (Invitrogen, Thermo Fisher Scientific) for 48 hours at room temperature with mild rotation, replacing with fresh EDTA after 24 hours. After demineralisation the bone pellet was washed three times with 100 µL 50 mM Tris solution (Invitrogen, Thermo Fisher Scientific). Each wash was added to the EDTA supernatant fractions. For buffer exchange the two EDTA supernatant fractions were passed through a 3 kDa molecular weight cut-off filter (Amicon, Sigma-Aldrich). An extraction buffer consisting of 2 M guanidine hydrochloride (Sigma-Aldrich), 10 mM Tris(2-carboxyethyl)phosphine (TCEP; Sigma-Aldrich), 20 mM 2- Chloroacetamide (CAA; Sigma-Aldrich), and 100 mM Tris was added to the proteins from the EDTA supernatant fractions to a final volume of 300 µL. The bone pellet was suspended in 300 µL of the same extraction buffer before incubation, along with the supernatant fraction, at 80°C under mild agitation for 2 hours. After incubation the protein concentration was measured for both the resuspended pellet and supernatant fraction using a Bicinchoninic acid assay (BCA) using a Pierce BCA Protein Assay kit (Thermo Scientific). For each sample, 50 µg proteins were aliquoted to a clean microtube. Sequential digestion was performed by first adding 1 µL 0.2 µg/µL LysC (Promega, VA1180) and incubating at 37°C under mild agitation for 2 hours, and then diluting to 0.6 M using a 25 mM solution consisting of 0.1 M Tris and 10% Acetonitrile (ACN; Thermo Fisher Scientific), before adding 1 µL 0.8 µg sequence grade Trypsin (Promega, V5111) and incubating overnight at 37°C. After approximately 18 hours the digestion was stopped by acidifying each sample to pH 2 using 10% trifluoroacetic acid (TFA; Sigma-Aldrich). A total of 10 µg protein (5 µg from pellet and supernatant fraction each) were then cleaned and desalted using EvoTip C18 tips (EV-2011, EvoSep).

#### 2.2.2. LC-MS/MS methods

Liquid chromatography tandem-mass spectrometry (LC-MS/MS) was performed on all samples using an EvoSep One (EvoSep) coupled to an Orbitrap Exploris 480 mass spectrometer (Thermo Fisher Scientific). On the EvoSep One were the peptides separated with the 100 samples-per-day (SPD) method with a 11.5 minute gradient.

The gradient used two mobile phases at a flow rate of 1.5 µL/minute. The two phases are: A) 5% acetonitrile (LC-MS grade ACN, VWR), 95% deionised water, and 0.1% formic acid (FA; LC-MS grade FA, Thermo Fisher Scientific), and B) 100% ACN and 0.1% FA. For the chromatography system an in-house made column (15 cm long silica tube, with inner diameter of 150 µm, packet with reprosil C18 particles of 1.9 µm diameter and 120 Å pores (ReproSil-Pur, C18-AQ, Dr. Maisch)) was attached to an easySpray source with a column oven at 60°C, source voltage of +2,000 V, and an ion transfer tube maintained at 275 °C.

The Orbitrap Exploris 480 mass spectrometer was operated in data-dependent acquisition (DDA) mode with the first MS1 scan set to a resolution of 60,000 within the mass range of 350-1,400 m/z. The top 12 ions selected for fragmentation had a minimum intensity above 2^e5^ and a charge state in the range of two to six. MS2 scans were acquired using higher-energy collision dissociation (HCD) with a resolution of 15,000, a 30% normalised collision energy, and an isolation width of 1.3 m/z.

### 2.3. Data analysis

All raw data (.raw format) were analysed using the software MaxQuant (version 2.1.3.0) (Cox & Mann, 2008). The main database search was conducted using a reference database consisting of canonical sequences from the whole human reference proteome (UP000005640; 83,413 entries; downloaded from Uniprot.org on June 3rd 2024). The search was run in specific Trypsin/P mode with Carbamidomethyl (C) as fixed modification, and Deamidation (NQ), Oxidation (MP), Gln (Q) → pyro-Glu, and Glu(E) → pyro-Glu as variable modifications. Other settings were set as default. To detect potential contamination the internal MaxQuant contaminant list was used. In addition, a search in MaxQuant was performed against the SPIN protein database (Rüther et al., 2022), utilising the same search parameters as the main database search.

The ten laboratory blanks were analysed in the same manner to determine any potential laboratory and cross sample contamination. Low levels of collagen type 1 were detected in the laboratory blanks, most likely indicating minimal sample carryover during the mass spectrometry analysis. All laboratory blanks were excluded from further analysis. Protein groups within the bone samples that matched only to the internal MaxQuant contaminant list, as well as all forms of keratin, were excluded from data analysis.

Mass spectrometry proteomic data, in .raw format, as well as MaxQuant output and SPIN analysis, have been deposited to the ProteomeXchange Consortium via the PRIDE (Perez-Riverol et al., 2022) partner repository with the dataset identifier PXD061703.

STRING analysis (Szklarczyk et al., 2025) was performed to gain insight into functions of the proteins identified, based on Gene Ontology (GO) terms. For analysis of cartilage-related proteins, the following GO terms were used: “Chondrocyte differentiation” (GO:0002062), “Cartilage condensation” (GO:0001502), and “Chondrocyte development” (GO:0002063). For analysis of immune-related proteins the following GO terms were used: “Humoral immune response” (GO:0006959), “Antimicrobial humoral immune response mediated by antimicrobial peptides” (GO:0061844), “Antimicrobial humoral response” (GO:0019730), “Inflammatory response” (GO:0006954), “Immune effector process” (GO:0002252), “Immune response” (GO:0006955).

All data analysis was performed in R (version 4.2.2) (R Core Team, 2022), utilising Rstudio (version 2024.12.0+467) (RStudio Team, 2022) and the following packages: *Janitor* (version 2.2.0) (Firke, 2023), *Tidyverse* (version 2.0.0) (Wickham et al., 2019), *ggpubr* (version 0.6.0) (Kassambara, 2023), *Patchwork* (version 1.3.0) (Pedersen, 2024), *Paletteer* (version 1.6.0) (Hvitfeldt, 2021), *Proteus* (version 0.2.16) (Gierlinski et al., 2018), *UpSetR* (version 1.4.0) (Conway et al., 2017) and *pheatmap* (version 1.0.12) (Kolde, 2019). Statistical tests were performed using the following packages: *Stats* (version 4.2.2) (R Core Team, 2022) and *FSA* (version 0.9.5) (Ogle et al., 2023). Additionally, the MaxQuant output was run through the Species by Proteome INvestigation (SPIN) computational pipeline for quality control (Rüther et al., 2022). R code for all analyses are available on Zenodo with the dataset identifier https://doi.org/10.5281/zenodo.15489291. Deamidation rates were calculated according to Mackie et al. (M. Mackie et al., 2018).

## 3. Results

We performed a quality control based on MS/MS data acquisition and PSM identification, as well as including summary parameters from the SPIN data analysis pipeline (Rüther et al., 2022). We observe full separation between laboratory blanks and bone samples when comparing the number of MS/MS scans acquired, relative protease intensities, and SPIN amino acid site counts (SI Figure 1). For the bone samples, the percentage of MS/MS identified ranges between 7 and 17%, which is on the high side for palaeoproteomic datasets (Chiang et al., 2024). From this we conclude that in general our specimens are well preserved, as also suggested by their chronological age and the preservation conditions of the Saint Eusebius church graveyard, and that the data analysis has generally saturated the possible MS2 identifications.

**Figure 1.**
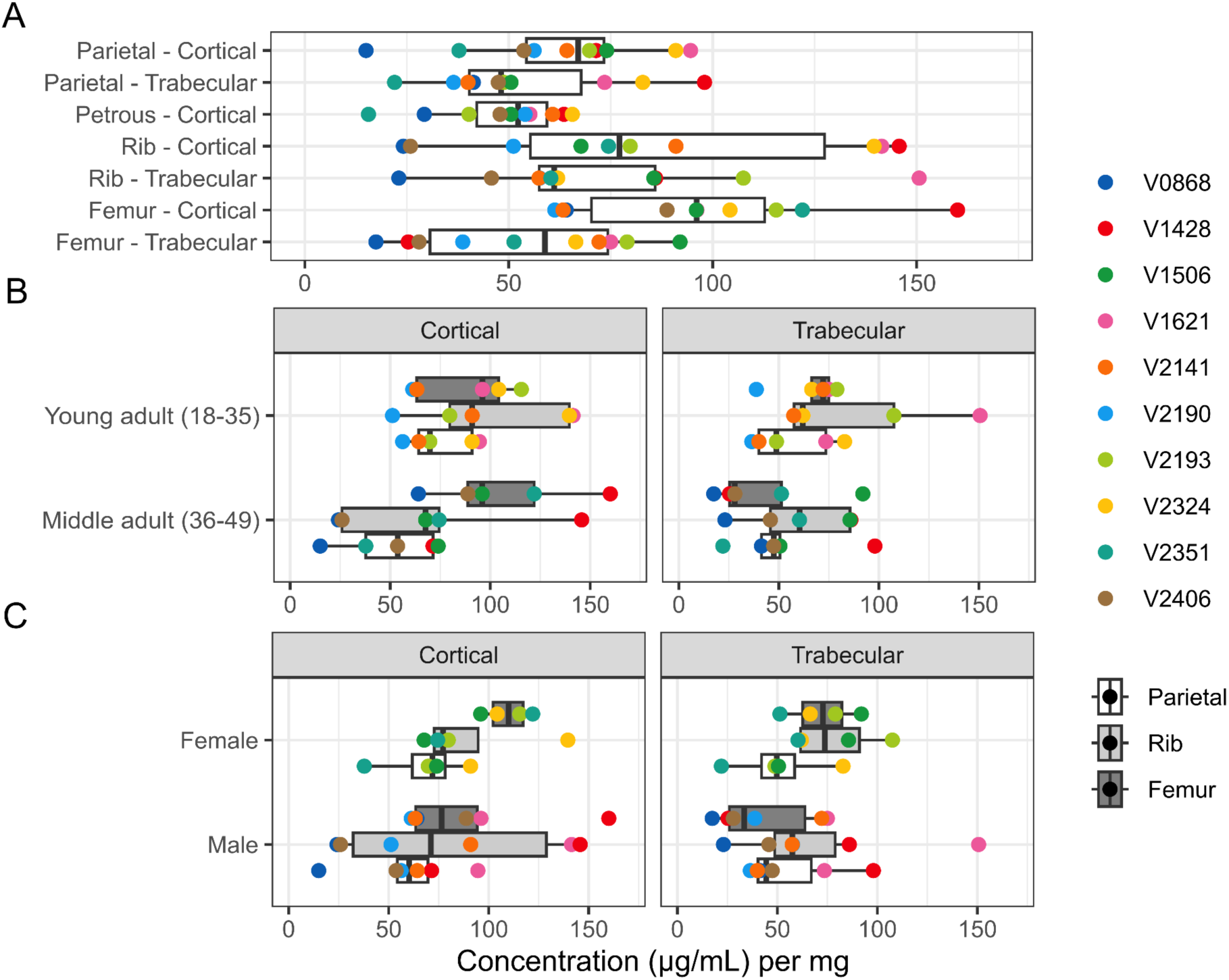
Bone protein concentrations across the archaeological human skeleton. A) Overall bone protein concentrations. B) Bone protein concentrations in relation to biological age, separated by cortical and trabecular bone samples. C) Bone protein concentrations in relation to biological sex, separated by cortical and trabecular bone samples. In each case, protein concentrations have been adjusted by the amount of bone powder sampled.

The bone samples all fall within range of each other when it comes to the SPIN site count and relative protease intensity, with the exceptions of the femur growth plate samples from V0868, which has higher relative protease intensity compared to all other bone samples, and from V2351, which has a lower site count than other bone samples. The remnants of the fused femoral growth plate proved to be a challenging location to sample, as its exact location is not visible to the naked eye. Due to the challenging sampling nature and inconsistency within the site counts and relative protease intensities, the femoral growth plate proteome is excluded from further analysis. Additionally, the parietal suture samples as well as the sample from the interior of the parietal bone are also excluded from further analysis due to similarities with the extracted external parietal bone proteome.

### 3.1 Protein concentration

We quantified protein concentration after bone demineralisation and solubilization. Controlling for the amount of bone sampled, and taking into account the cortical-trabecular bone pairs for the parietal bone, rib, and femur, we observe slightly higher protein concentration in cortical bone compared to trabecular bone across our three cortical-trabecular bone pairs (Paired t-test, p = 0.004; Figure 1). We observe no significant difference between protein concentration in the cortical bone of bone elements formed through endochondral ossification compared to the cortical bone of bone elements formed through intramembranous ossification (Welch two sample t-test, n.s.). In this context, it is noteworthy that the protein concentration observed for the petrous bone, formed through endochondral ossification, is lower than that observed femur and rib cortical bone, and comparable to the parietal cortical bone. The four locations sampled across the parietal, a suture, trabecular bone, and internal and external cortical bone, all have rather similar protein concentrations. Furthermore, we observe no significant relationship between protein concentrations of cortical-trabecular bone pairs for either biological sex or age (Welch two sample t-test, n.s.).

### 3.2 Proteome size & composition

Based on differences in protein concentration and previous research indicating compositional differences in archaeological cortical and trabecular bone proteomes, we first compared proteome composition between these two tissue types in each of three bone element pairs (femur, rib, parietal). We find that trabecular bone proteomes are, in comparison to their corresponding cortical bone proteomes, smaller in size (Paired t-test, p < 0.001) and have lower peptide counts associated with them (Wilcoxon signed rank test, p < 0.001; Figure 2, SI Figure 2). Although the trabecular bone proteomes are smaller than their cortical counterparts, there are some proteins only identified in trabecular bone proteomes, such as cartilage intermediate layer protein 2 (CILP2; see below). In contrast, matrix metalloproteinase-2 (MMP2; also known as 72 kDa type IV collagenase), transmembrane protein 119 (TMEM-119), olfactomedin-like protein 1 (OLFL1), adipocyte enhancer-binding protein 1 (AEBP1), and Kazal-type serine protease inhibitor domain-containing protein 1 (KAZALD1) are all only identified in cortical bone. These are all proteins with relevant roles during ossification (Bernardo et al., 2011; Hisa et al., 2011; Inoue et al., 2006; Jin et al., 2024; Kawao et al., 2024; Mosig & Martignetti, 2013; Murakami et al., 2018; Nyman et al., 2011; Sauer et al., 2022; Shibata et al., 2004; Tanaka et al., 2012).

**Figure 2.**
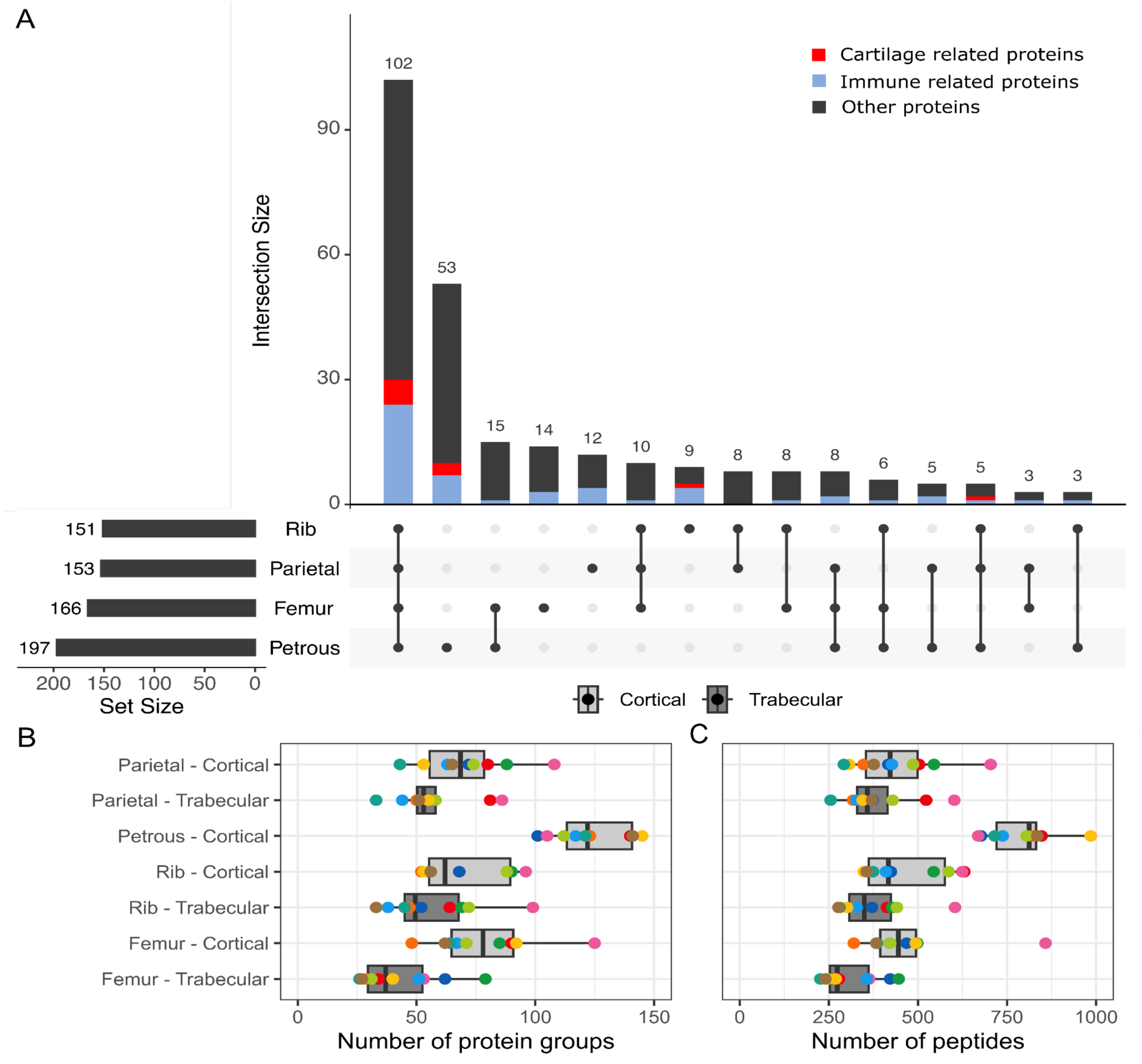
Proteome size and composition. A) Proteome size and shared proteins across cortical bones. B) Proteome size. C) Peptide counts. Specimen colours in panels B and C are the same as in Figure 1.

Our dataset includes sampling locations initially ossified through endochondral ossification (femur, rib, petrous bone) and intramembranous ossification (parietal bone). Considering only cortical bone sample locations, we observe differences in terms of proteome size when comparing the number of identified protein groups (Kuskal-Wallis test, p < 0.001) and the number of peptides (Kruskal-Wallis test, p < 0.001). The petrous pyramid stands out for both observations, with high protein counts as well as larger peptide numbers (post-hoc Dunn’s test, all p < 0.05; SI Figure 4). There are no significant differences in proteome size or peptide counts for the other cortical locations (post-hoc Dunn’s test, n.s.; SI Figure 4).

When comparing protein abundance of the cortical bone samples we find that, when excluding the petrous bone, osteocalcin (OSTCN) is more abundant in femoral and rib samples compared to the parietal bone samples (Figure 3). Osteocalcin is a non-collagenous protein that is mainly secreted by osteoblasts. Smaller amounts are, however, secreted by hypertrophic chondrocytes (Zoch et al., 2016). Osteocalcin has a role in bone mineralisation and it, alongside osteopontin, forms a scaffold with collagenous proteins for the mineral deposition in the bone (Sroga et al., 2011). Osteocalcin has also been shown to be in higher abundance in newer bone tissue, suggesting its involvement in bone remodeling and metabolism. The exact role of osteocalcin in bone remodeling and metabolism is currently not fully understood (Sroga et al., 2011; Zoch et al., 2016).

**Figure 3.**
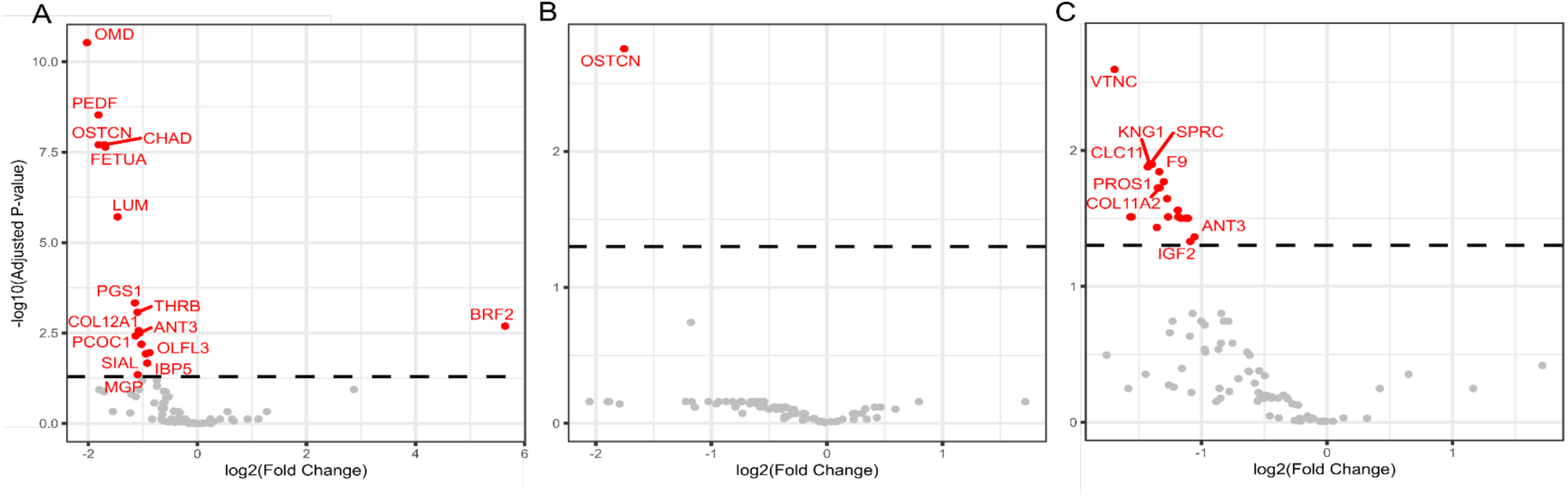
Protein abundance differences. A) Based on bone type. Cortical and trabecular bone pairs of femur, rib, and parietal bone. A negative fold change indicates higher abundance in cortical bone, and a positive fold change indicates higher abundance in trabecular bone. B) Based on ossification processes. Cortical bone proteomes of femur, rib, and parietal bone. C). Based on ossification processes of cortical bone proteomes of petrous, femur, rib, and parietal bone. For panels B and C a negative fold change indicates higher abundance in endochondral ossification and a positive fold change indicates higher abundance in intramembranous ossification. Dashed line in all panels indicates the used threshold of significance (p < 0.05).

**Figure 4.**
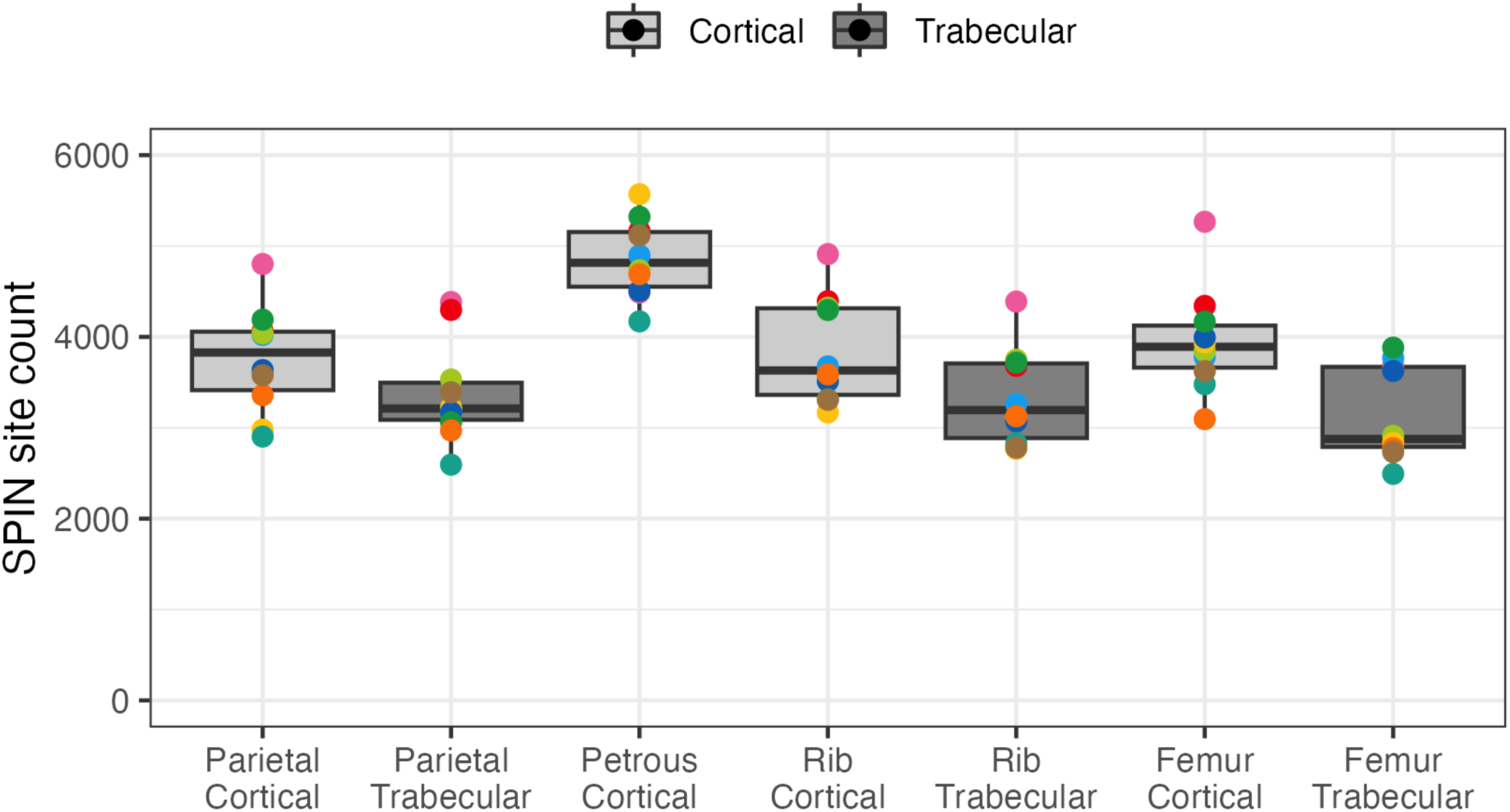
Amino acid site counts identified through SPIN analysis, per sampling location. Specimen colours are the same as in Figure 1.

When including the petrous bone, several proteins (such as vitronectin (VTNC), alpha- 2-HS-glycoprotein (FETUA), and osteomodulin (OMD), among others) have higher abundance in the petrous bone samples compared to other cortical bones (Figure 3). The larger size of the petrous bone proteome, the higher peptide counts, the comparatively low protein concentration, and its enhanced degradation (see below) all suggest that, for the non-petrous bone samples initially formed through endochondral ossification, differences in initial bone ossification have largely been obscured by subsequent bone remodelling during the adult life of the studied individuals.

Endochondral ossification is directly associated with the extracellular expression of a unique set of proteins by (hypertrophic) chondrocytes. Thirteen cartilage related proteins were identified based on GO terms (see 2.3. Data analysis; Figure 2). Four of these proteins are only secreted by chondrocytes, and not osteoblasts, and are therefore considered to be cartilage-specific. These proteins are collagen type X (Alini et al., 1996; Deng et al., 2018), aggrecan core protein (Wilson et al., 2008), cartilage intermediate layer protein 2 (CILP2) (Bernardo et al., 2011), and cartilage matrix protein (MATN1) (Y. Chen et al., 2016; Pei et al., 2008). These proteins are all expressed during initial ossification. In our dataset, these proteins are only present in adult bone samples formed through endochondral ossification. ACAN and CLP2 have the highest number of unique peptides in femoral trabecular bone. COL10A1 is most abundant in the petrous bone, while lower numbers are observed in femoral and rib trabecular bones. MATN1 is only present in the petrous bone (SI Figure 5).

**Figure 5.**
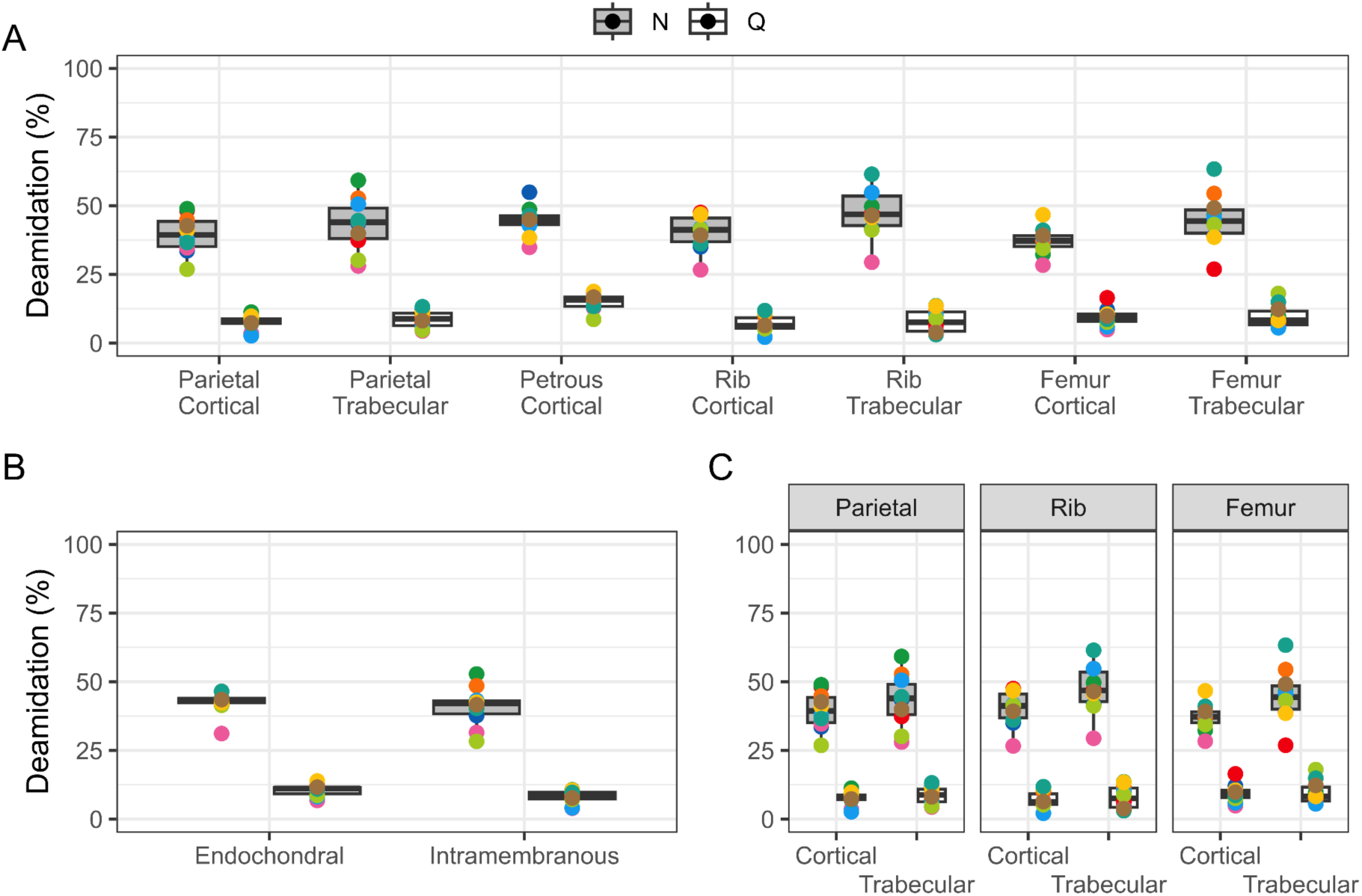
Deamidation of asparagine (N) and glutamine (Q). A) Overall deamidation. B) Deamidation of the two ossification processes for all sample locations. C) Deamidation of cortical and trabecular bone pairs from femur, rib, and parietal bone. For all panels: 0% deamidation indicates no N or Q deamidation, respectively, while 100% indicated full deamidation. Colour of points indicating individuals are the same as in Figure 1.

The bone proteomes recovered also contain information derived from the individual’s immune system, based on the GO terms, where 40 proteins were identified. Additionally, we identified three immunoglobulin proteins (Immunoglobulin kappa constant (IGKC), Immunoglobulin heavy constant gamma 1 (IGHG1), and Immunoglobulin heavy constant mu (IGHM)) that are not included in the GO terms. Immunoglobulins have been proposed to study the health of past populations through their presence in dental enamel (Buonasera et al., 2024; Wilkin et al., 2024). Of these immunoglobulins, IGHG1 is present in samples from all individuals. IGKC and IGHG1 are identified in all skeletal elements, while interestingly, IGHM is not identified in the petrous. Out of a total of 43 immune related proteins identified, 29 were observed to have the highest number of peptides within the petrous bone, and additional 6 proteins were only found in the petrous bone (Figure 2). Most immune related proteins are identified in all skeletal locations (Figure 2), but there are also immune-related proteins only identified in the femur, the rib, or the parietal. Together, this indicates that accessing health-related palaeoproteomic information might be dependent on, or restricted to, protein presence in particular skeletal locations.

The petrous bone also stands out when comparing the whole bone proteome per individual in hierarchical clustering and falls as an outgroup for nine out of the ten individuals. For seven out of ten individuals, clustering of cortical and trabecular bone is additionally present, suggesting that proteome variation is largely driven by bone type and its maintenance (SI Figure 3).

Ultimately, the phylogenetic power of a palaeoproteomic study is dependent on the maximization of the number of amino acid positions that can be called confidently for as many different proteins as possible. Sequence coverage, here reported as site counts based on SPIN analysis, varies based on bone and bone type. We observe significantly higher sequence coverage in cortical bone proteomes compared to trabecular bone proteomes (Two-way ANOVA, F = 38.48, p < 0.001; Figure 4). Additionally, which skeletal element is sampled effects the sequence coverage too (Two-way ANOVA, F = 8.73, p < 0.001; Figure 4), where the highest sequence coverage is observed in the petrous bone (Tukey’s HSD, p < 0.001 in each case; SI Figure 4). Generally speaking, cortical bone is therefore more suited for conducting palaeoproteomic phylogenetic studies than trabecular bone. Among cortical bone types, the petrous stands out as particularly promising for phylogenetic studies. This latter observation is noteworthy, since the studied individuals are only a couple of centuries old, and this difference in protein sequence coverage for the 20 proteins considered in SPIN is already apparent.

### 3.3 Deamidation

Deamidation, a diagenetic modification of the amino acids asparagine (N) and glutamine (Q), has been applied in various palaeoproteomic context to quantify degradation of proteins over time (Ásmundsdóttir et al., 2024; F. Chen et al., 2019; M. Mackie et al., 2018; Nair et al., 2023; Ramsøe et al., 2020; Schroeter & Cleland, 2016; Welker et al., 2016). We calculated deamidation values of asparagine and glutamine for all proteins across the dataset, and filtered out proteins with fewer than five peptides across the ten individuals. We observe overall higher deamidation of asparagine compared to glutamine (Wilcoxon rank sum exact test, p < 0.001; Figure 5). This is in agreement with previous theoretical and experimental research, including palaeoproteomics studies (Mylopotamitaki et al., 2023; Ramsøe et al., 2020; Robinson & Robinson, 2001).

Deamidation values differ between bone pairs with both cortical and trabecular bone (femur, rib, and parietal) for asparagine deamidation, where we observe a higher rate in trabecular bone compared to cortical bone (Paired t-test, p < 0.001; Figure 5). For this faster- deamidating amino acid, the observation therefore mirrors what was observed for cortical/trabecular bone pairs in early Holocene specimens from Spain (Ásmundsdóttir et al., 2024). This is not observed for glutamine deamidation, where the rate for the three pairs is similar for both bone types (Paired t-test n.s.; Figure 5).

We observe a higher rate of glutamine deamidation for bones formed through endochondral ossification compared to intramembranous ossification (Welch two sample t-test, p = 0.02; Figure 5), this is, however, not observed in asparagine deamidation (Welch two sample t-test, n.s.). When it comes to either biological age or biological sex, we observe no differences in either absolute asparagine or glutamine deamidation (Welch two sample t-Test, n.s.). How deamidation rates differ among and between other age groups, for example children and adolescents, is currently unknown.

Restricting our analysis to cortical bone samples, we observe differences in glutamine deamidation rates across all locations (One-way ANOVA, F = 14.84, p < 0.001), where the petrous pyramid notably stands out from the parietal, rib and femoral cortical bone samples with higher glutamine deamidation rates (Tukey’s HSD, p < 0.05 in all cases; SI Figure 4). We find it unlikely that this is the result due to the higher bone density or lower water content of the petrous bone, both of which would be expected to result in lower extents of glutatmine deamidation. Rather, we suggest that this is the result of extended exposure of the petrous bone proteome to body temperatures, while undergoing no to minimal bone turnover during life. Future research including younger and older age cohorts can confirm that the extent of deamidation of the petrous bone proteome is correlated with biological age.

## 4. Discussion

The use of palaeoproteomic methods to study human evolution, by using either faunal or human remains, has greatly increased over the past few years. Increasing interest in this biomolecular approach has the potential risk of oversampling a limited archaeological resource since ancient biomolecular research is, by definition, destructive. Most fields within biomolecular archaeology, outside of palaeoproteomics, have optimised their sample selection strategies to improve their success rate, such as sampling the petrous bone for ancient DNA (Alberti et al., 2018; Damgaard et al., 2015; Hansen et al., 2017; Parker et al., 2020; Pinhasi et al., 2015) and sampling cortical bone for isotopic analysis (R. Hedges et al., 2008; R. E. M. Hedges et al., 2007). Here we present our results of proteome variations within the human skeleton with the aim of starting the process of optimising sample selection strategies for the field of palaeoproteomics.

### 4.1. Sampling selection strategies

What makes optimising palaeoproteomic sample selection strategies especially challenging is combining the influence of biology of the living bone with protein preservation through time. The bones in the human skeleton are formed in two different ways, endochondral ossification that utilises a cartilage template to form the bone, and intramembranous ossification which does not utilise the cartilage template. In our data we observe some of the cartilage-specific proteins, such as cartilage intermediate layer protein 2 (CILP2) and aggrecan core protein (ACAN), only in bones formed through endochondral ossification. We identify no proteins unique to intramembranous ossification. The presence of cartilage-specific proteins in adult bone is noteworthy, since these proteins derive from steps in the ossification process that were completed years, if not decades, before the death of these individuals. As a result, subsequent bone remodeling would have been expected to remove them. Their continuing presence in some of the specimens sampled here indicates that bone remodeling had not removed all bone initially ossified yet.

Most bones in the human body are composed of two bone types, cortical and trabecular bone, that differ in terms of bone density and water content (Beresheim et al., 2020; Faraldi et al., 2022; Gong et al., 1964; Haverfield et al., 2023). In cortical and trabecular bone pairs from the femur, rib, and parietal bone, we observe larger and more complex proteomes within the cortical bone samples. Additionally, these two bone types have different rates of turnover, with faster rates in trabecular bone (Deftos, 1998; Parfitt, 1994, 2002; Parfitt et al., 1996). We observe the most variation between the two different bone types when it comes to proteome size, both in the number of protein groups and the number of peptides. Two skeletal locations, the fused growth plate of the distal femur and cranial sutures of the parietal bone, were included in this study as they stood out from other locations in terms of formation and potential proteome composition. The distal femoral growth plate is the last place to ossify in the femur, and in theory could contain more cartilage-specific proteins than femoral cortical or trabecular bone. Within our data, there are minimal differences between the growth plate and the femoral trabecular bone samples. The growth plate is very difficult to precisely locate, however, as it is not visible externally in adult bone. More precise sampling, for example with the help of X-rays, of the proximal tibial growth plate, which is sometimes visible on the bone surfaces of adult human skeletons,into adult life could shed more light onto the possible composition of the growth plate proteome.

The other location, cranial sutures of the parietal bone, are fully ossified and fuse relatively late in life, between the ages of 25 and 30 (Celik et al., 2021). With this late fusion there is less time for loss of proteome variation through turnover compared to other cortical bone locations. Unfused cranial sutures have been observed to have cartilage related proteins, such as COL2A1, COL10A1, and aggrecan core protein, that are normally not observed in cranial bones formed through intramembranous ossification (Coussens et al., 2007). We do not observe any cartilage related proteins within our cranial suture specimens, nor any proteins uniquely observed within the sutures. This is most likely due to the biological age of the individuals in this study. Sampling cranial sutures in younger individuals, especially children and adolescents, might reveal the presence of such these proteins.

The petrous pyramid is one of the densest bones in the human body and undergoes minimal to no turnover during life (Harvig et al., 2014; Jørkov et al., 2009). In our data, the petrous bone does not stand out when it comes to protein concentration. It does, however, stand out when it comes to proteome size and composition, as well as in glutamine deamidation. The petrous proteome is larger than other cortical bone samples, and contains proteins, albeit in low concentrations, that are vital during the initial ossification, such as cartilage matrix protein (MATN1) and collagen type X (COL10A1). This is direct evidence for the preservation of proteins excreted by (hypertrophic) chondrocytes during initial endochondral bone ossification. Furthermore, this observation in our petrous bone samples indicates that throughout the individual’s life a large portion of the proteome is lost due to bone turnover in the other bone elements. How fast these proteins are lost, and the influence of protein preservation on this observation, is currently not well understood. The presence of various proteins either only observed in the petrous bone, or observed there in higher abundance, the overall larger proteome size, as well as the significantly higher reconstructed site count, indicates that the petrous bone is preferred over other skeletal locations when it comes to sample selection. Our observations are made on archaeological human bone proteomes that are only a couple of centuries old. Bone mineral density and water content are known to determine the rate and fate of deep-time protein survival in mineralised tissues, including hominin skeletal material (Cappellini et al., 2019; Demarchi et al., 2016, 2022; Welker et al., 2020). As a result, over the long time spans during which proteomes are preserved in Pleistocene hominin materials, the effect of a higher bone mineral density and a lower water content in petrous bones can be expected to, relatively speaking, make the petrous bone stand out even more compared to other bone tissues. Both for recent archaeological and deep-time palaeoproteomics research with a strong phylogenetic focus, the petrous bone is therefore the optimal source of a large, complex ancient proteome.

This, however, places an increased sampling pressure on the petrous bone as other biomolecular fields studying human evolution, such as ancient DNA and isotopic research, prefer the petrous bone as well. In addition, sampling the petrous bone is always (highly) destructive (Charlton et al., 2019; Källén et al., 2024; Pinhasi et al., 2015, 2019). With only two specimens available within a skeleton, that is *if* both are preserved, sampling petrous bones for palaeoproteomics research limits reproducibility and/or revisiting specimens with new or improved analytical methods in the future. We therefore advocate for conservative considerations when proposing the sampling of petrous bones for (deep-time) hominin phyloproteomics.

We observe both higher protein concentration and larger proteomes in bones formed through endochondral ossification, especially cortical bone, compared to bones formed through intramembranous ossification. We therefore recommend sampling cortical bone, especially when specimens are suspected to have poor preservation (Ásmundsdóttir et al., 2024). We note, however, that some proteins known to contain phylogenetically informative positions among hominin populations, such as COL1A2, COL2A1, COL10A1 and MGP, have a higher abundance the petrous and trabecular bone of the femur and rib. When the petrous is not available, or its sampling not advisable, there might therefore be an interest in sampling both cortical and trabecular bone of skeletal elements forming through endochondral ossification. Together, this would maximise the possibility of recovering phylogenetically informative positions in a range of proteins.

### 4.3. Deamidation

Deamidation is one of the main methods used to estimate protein degradation and thus authenticity of ancient proteins. Over time, the amino acids asparagine (N) and glutamine (Q) are transformed to aspartic acid and glutamic acid, respectively. The rate at which these two amino acids are transformed differs between the two, with asparagine deamidation being faster than glutamine deamidation (Ásmundsdóttir et al., 2024; M. Mackie et al., 2018; Nair et al., 2023; Ramsøe et al., 2020; Robinson et al., 2004; Wright, 1991). Various factors are involved in the rate of deamidation other than time, such as temperature and soil pH (Kendall et al., 2018). Minimal levels of deamidation are also observed during an individual’s life (Boudier- Lemosquet et al., 2022).

We observe higher rate of glutamine deamidation in bone formed through endochondral ossification compared to bone formed through intramembranous ossification, indicating that the initial mode of ossification does influence the rate of deamidation observed in the adult, archaeological human skeleton. For the two bone types, cortical and trabecular bone, we observe a higher rate of asparagine deamidation in trabecular bone compared to cortical bone. The higher rate of deamidation in trabecular bone is most likely due to the properties of the bone, such as a lower mineral density as well as a higher water content, resulting in faster degradation.

The petrous bone stands out in terms of glutamine deamidation, with a higher rate of glutamine deamidation compared to all other skeletal locations sampled. The higher mineral density and the lower water content of the petrous bone can be expected to result in lower rates of diagenetic protein modification (Demarchi et al., 2016; Hansen et al., 2017; Kontopoulos et al., 2019), and can therefore not explain our experimental observations. Instead, we suggest that this enhanced glutamine deamidation rate is due to the minimal bone remodelling of the petrous bone during life. As a result, the proteins embedded in the petrous bone have been exposed to body temperatures for longer periods of time, compared to continuously replaced bone proteins in other skeletal locations, resulting in enhanced deamidation rates of the proteins preserved in this unique biomineral location. In a given preservation environment, glutamine deamidation of petrous bone proteins would therefore correlate with biological age, something that can be tested by studying younger and older age cohorts. If verified, this would provide a relative approach of biological age estimation that would be of use in forensics and osteoarchaeology (Baker, 2016; Clark et al., 2020, 2023; Mahlke et al., 2021).

## 5. Conclusions

Here we combined bone biology with the archaeological preservation of proteins to gain a deeper understanding of the bone proteome composition, and its heterogeneity, in archaeological skeletal elements. In bones formed by different ossification processes, endochondral and intramembranous ossification, we observe minimal differences in protein concentrations. When it comes to proteome size, we observe larger proteomes in endochondral bone, especially in endochondral cortical bone. These endochondral bone samples also include a few cartilage-specific proteins, and these are absent in the intramembranous bone samples. The petrous bone stands out with the largest and most complex proteome. The composition of the petrous bone proteome suggests that during and right after ossification, bone proteomes in general are more complex, and that some of the variation is lost during the individual’s life through bone remodeling, which is minimal in the petrous bone compared to other locations.

Additionally, we observe that for the two bone types, cortical and trabecular bone, the proteomes of cortical bone are larger, in terms of concentration in the bone and in terms of proteome size, as well as less degraded than their trabecular bone counterparts. Trabecular bone, however, is observed to preserve a few phylogenetically informative proteins, such as COL2A1, at a higher concentration than cortical bone.

Our results indicate that the petrous bone is the ideal source to recover large, complex bone proteomes, already in skeletal material of only a couple of centuries old. Due to its higher mineral density and lower water content, across larger time depths, the petrous bone can be expected, compared to other cortical bone locations, to preserve the proteins contained within it even better. The petrous is therefore the ideal bone element location to conduct palaeoproteomic phylogenetic analysis. This might place an increased burden on the already limited number of petrous specimens available for archaeological science research. We acknowledge that petrous bone sampling is highly destructively, difficult to replicate, and will destroy a piece of the (hominin) skeleton informative on evolutionary processes through its morphology (Smith et al., 2024; Stoessel et al., 2016). Rather than competing for specimen access with other disciplines, such ancient DNA analysis and isotopic research, and to ensure preservation of its internal morphology, we recommend that palaeoproteomics researchers explore the utility of cortical bone sampling, potentially combined with trabecular bone sampling to increase proteome size and complexity. We show that cortical bone proteomes, especially those deriving from endochondral ossification, are generally larger and better preserved than trabecular ones, and contain a larger number of informative amino acid positions. Nevertheless, trabecular samples from endochondral bone locations might provide enhanced access to cartilage-related proteins, and these could be of specific phylogenetic interest, depending on the taxonomic context. Taken together, our research indicates that archaeological bone proteomes extracted from relatively recent, adult human skeletons reflect bone biology of the living skeleton. This knowledge should therefore guide future sampling strategies of archaeological and palaeontological materials, both for evolutionary and medical studies in palaeoproteomics.

## Acknowledgements

This research was supported by funding from the European Research Council (ERC) under the European Union’s Horizon 2020 research and innovation programme, grant agreement no. 948365 (PROSPER), awarded to F.W. R.D.Á. was supported by the Leakey Foundation. S.S. was funded by the Dutch Research Council (NWO) no. VI.Vidi.201.153). We thank Martin P. Defilet, senior management advisor for Heritage and Archaeology, and the city of Arnhem, the Netherlands, for access to skeletal remains. Mass spectrometry analysis, carried out at the Novo Nordisk Foundation Center for Protein Research, was funded in part by a donation from the Novo Nordisk Foundation (grant number NNF14CC0001). Additionally, we thank Dr. Felicia Fricke for osteological assistance in parietal bone orientation and femoral growth plate location, as well as Dr. Dorothea Mylopotamitaki and Anna-Lena Titze for assistance and instructions for petrous bone sampling.

