## Supplementary material for "Composition variations in archaeological human bone proteomes": SI Figure

### Supplementary Information

Ragnheiður Diljá Ásmundsdóttir[1], Gaudry Troché[1], Jesper V. Olsen[2], Sarah Schrader[3], Frido Welker[1]

1. Globe Institute, Faculty of Health and Medical Sciences, University of Copenhagen, Copenhagen, Denmark.
2. Novo Nordisk Foundation Center for Protein Research, Faculty of Health and Medical Sciences, University of Copenhagen, Copenhagen, Denmark.
3. Faculty of Archaeology, Leiden University, Leiden, the Netherlands.

SI Table 1: Sample weight (mg) per location for the ten individuals. Sutures were not present for parietal bone fragments of individuals V1506 and V2351.

| Bone | Type | **V0868** | **V1428** | **V1506** | **V1621** | **V2141** | **V2190** | **V2193** | **V2324** | **V2351** | **V2406** |
|---|---|---|---|---|---|---|---|---|---|---|---|
| **Parietal** | Cortical exterior | 20 | 20 | 21 | 19 | 19 | 19 | 19 | 19 | 20 | 20 |
| | Trabecular | 22 | 21 | 20 | 20 | 21 | 19 | 19 | 21 | 21 | 21 |
| | Cortical interior | 20 | 18 | 22 | 21 | 22 | 22 | 19 | 20 | 19 | 22 |
| | Suture | 21 | 20 | - | 19 | 20 | 21 | 21 | 22 | - | 19 |
| **Petrous** | | 21 | 22 | 19 | 20 | 20 | 22 | 24 | 22 | 20 | 22 |
| **Rib** | Cortical | 20 | 19 | 21 | 21 | 21 | 20 | 22 | 21 | 22 | 19 |
| | Trabecular | 22 | 21 | 19 | 19 | 22 | 20 | 19 | 21 | 19 | 20 |
| **Femur** | Cortical | 22 | 20 | 22 | 22 | 20 | 21 | 21 | 20 | 21 | 20 |
| | Trabecular | 21 | 20 | 21 | 19 | 21 | 21 | 21 | 21 | 21 | 21 |
| | Growth plate | 21 | 19 | 22 | 21 | 21 | 20 | 20 | 21 | 21 | 22 |

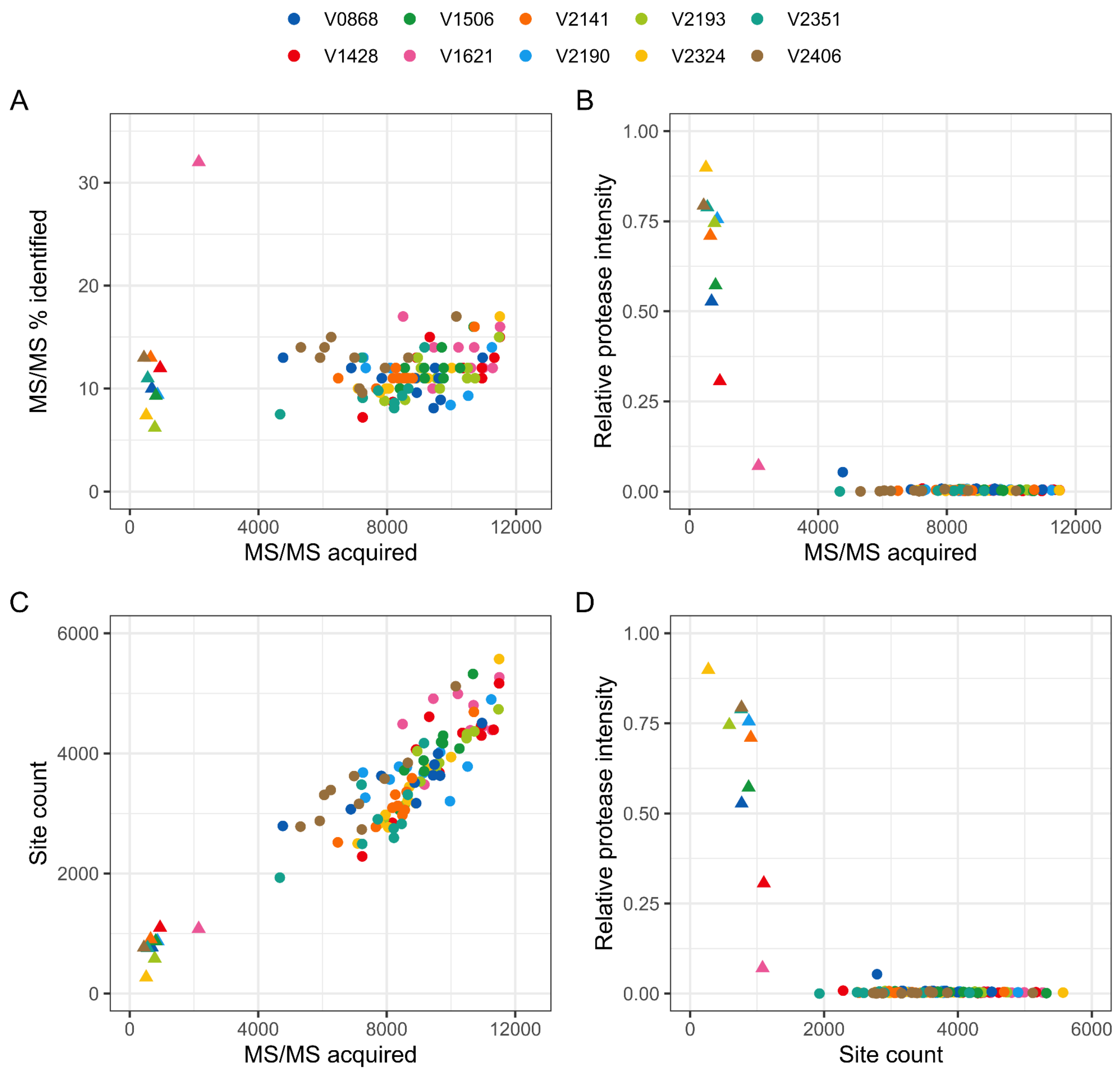


SI Figure 1: Quantitative comparisons of MS data acquisition and MS/MS PSM recovery between laboratory blanks (triangles) and bone samples (circles). A) MS/MS acquired and percentage of MS/MS identified. B) MS/MS acquired and relative protease intensity. C) MS/MS acquired and site count. D) Site count and relative protease intensity. Site count and relative protease intensity derive from the SPIN data analysis pipeline.

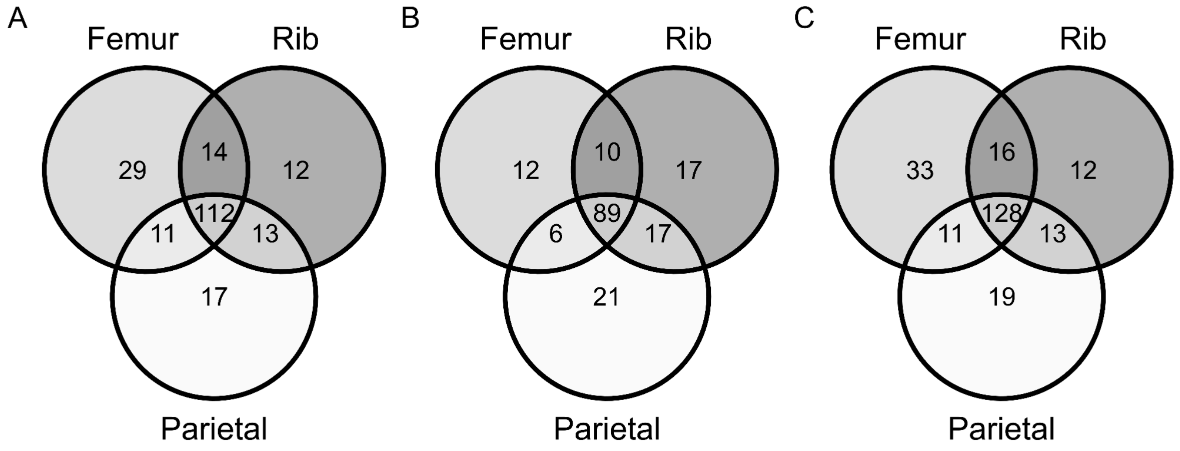


SI Figure 2: Unique and shared protein groups across femur, rib, and parietal bone. A) Cortical bone. B) Trabecular bone. C) Cortical and trabecular bone.

SI Figure 3: Hierarchical clustering of skeletal elements and protein groups per individual sampled. Clustering was performed using the Canberra method for non-parametric data using protein intensities with label free quantification (LFQ). Collagen type 1 (COL1A1 and COL1A2) are excluded due to their high abundance to prevent masking of proteins of lower abundance. This does not affect the final clustering.

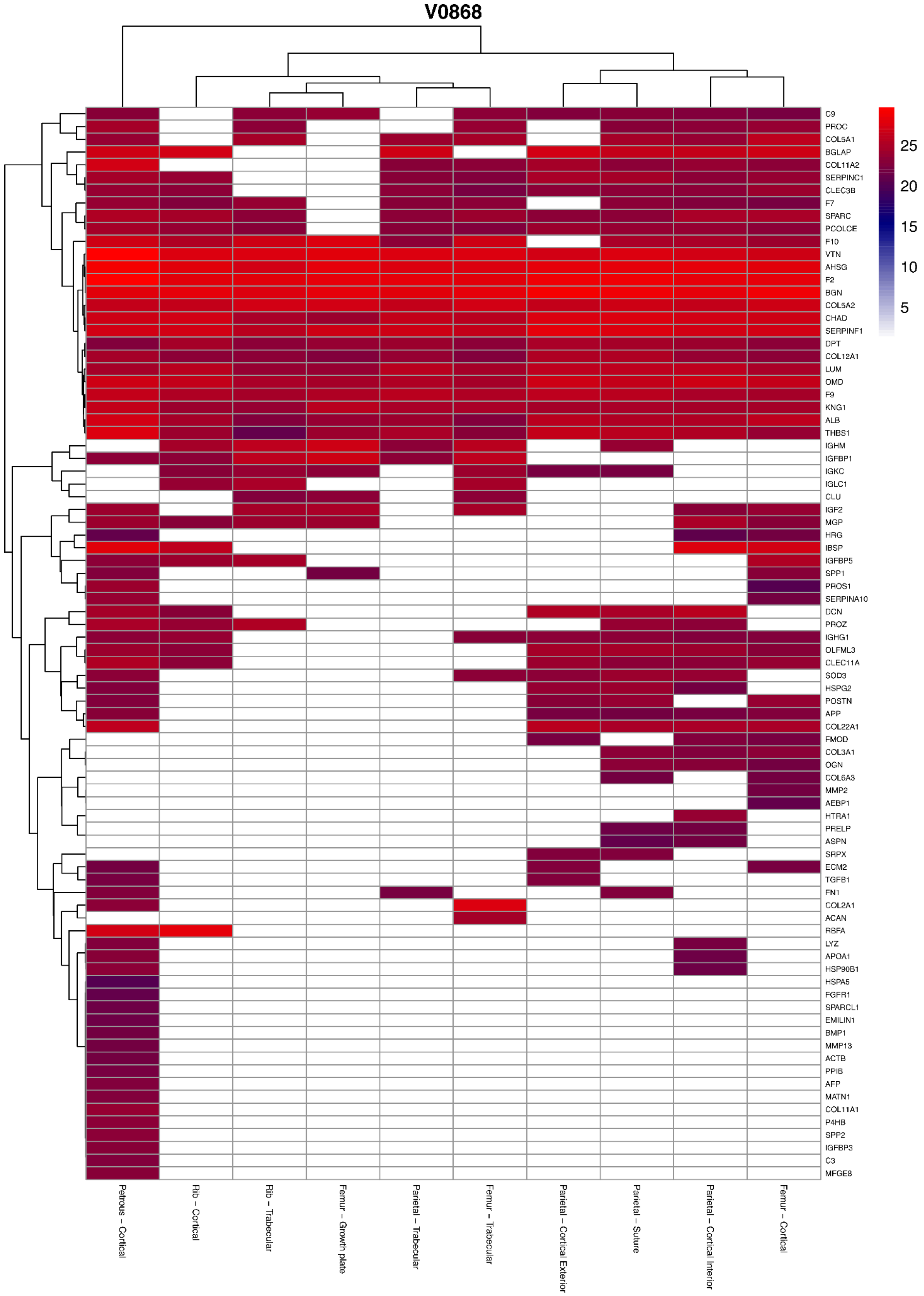
V0868
25
20
15
10
5
C9
PROC
COL5A1
BGLAP
COL11A2
SERPINC1
CLEC3B
F7
SPARC
PCOLCE
F10
VTN
AHSG
F2
BGN
COL5A2
CHAD
SERPINF1
DPT
COL12A1
LUM
OMD
F9
KNG1
ALB
THBS1
IGHM
IGFBP1
IGKC
IGLC1
CLU
IGF2
MGP
HRG
IBSP
IGFBP5
SPP1
PROS1
SERPINA10
DCN
PROZ
IGHG1
OLFML3
CLEC11A
SOD3
HSPG2
POSTN
APP
COL22A1
FMOD
COL3A1
OGN
COL6A3
MMP2
AEBP1
HTRA1
PRELP
ASPN
SRPX
ECM2
TGFB1
FN1
COL2A1
ACAN
RBFA
LYZ
APOA1
HSP90B1
HSPA5
FGFR1
SPARCL1
EMILIN1
BMP1
MMP13
ACTB
PPIB
AFP
MATN1
COL11A1
P4HB
SPP2
IGFBP3
C3
MFGE8
Petrous – Cortical
Rib – Cortical
Rib – Trabecular
Femur – Growth plate
Parietal – Trabecular
Femur – Trabecular
Parietal – Cortical Exterior
Parietal – Suture
Parietal – Cortical Interior
Femur – Cortical

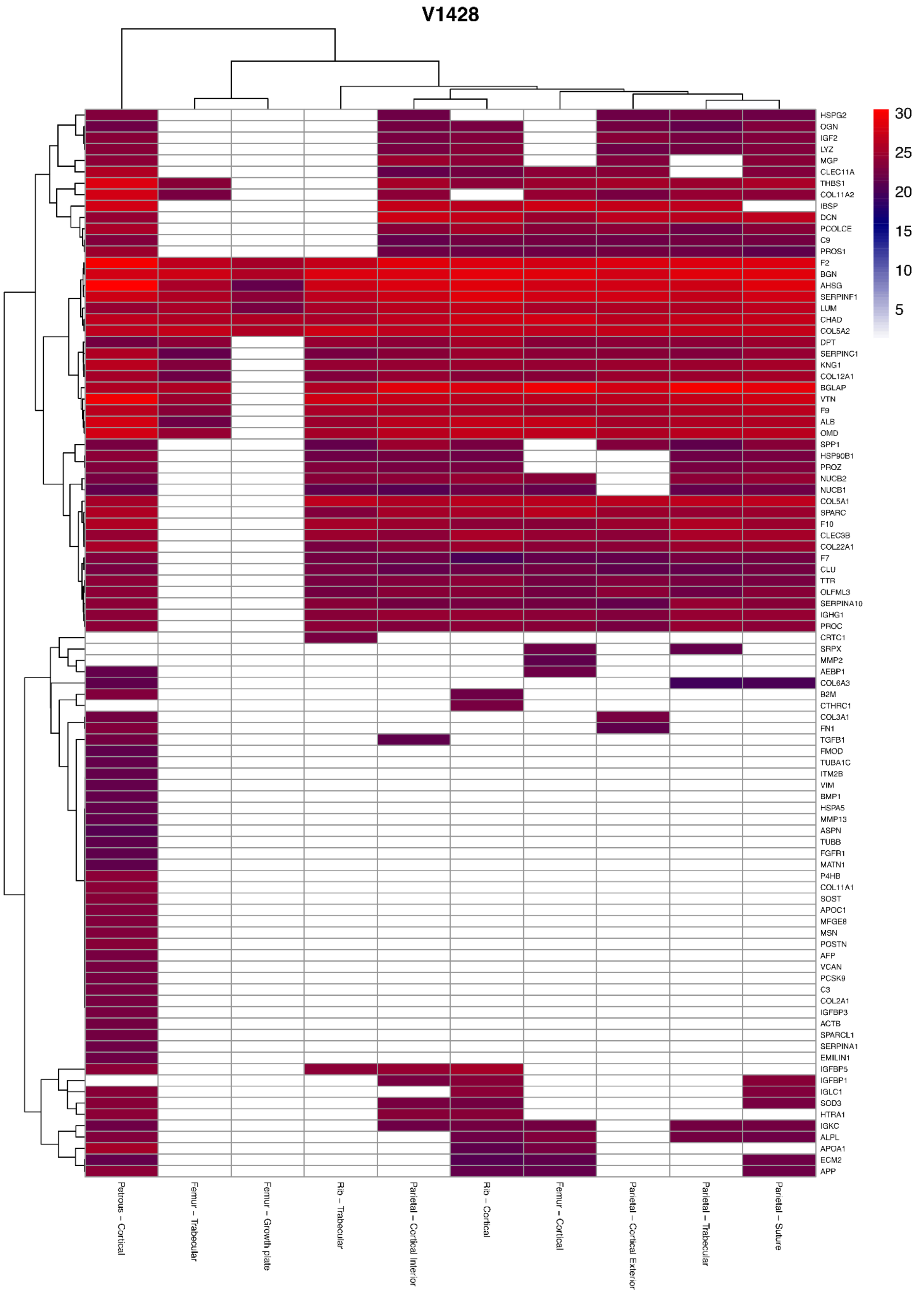
V1428
HSPG2
OGN
IGF2
LYZ
MGP
CLEC11A
THBS1
COL11A2
IBSP
DCN
PCOLCE
C9
PROS1
F2
BGN
AHSG
SERPINF1
LUM
CHAD
COL5A2
DPT
SERPINC1
KNG1
COL12A1
BGLAP
VTN
F9
ALB
OMD
SPP1
HSP90B1
PROZ
NUCB2
NUCB1
COL5A1
SPARC
F10
CLEC3B
COL22A1
F7
CLU
TTR
OLFML3
SERPINA10
IGHG1
PROC
CRTC1
SRPX
MMP2
AEBP1
COL6A3
B2M
CTHRC1
COL3A1
FN1
TGFB1
FMOD
TUBA1C
ITM2B
VIM
BMP1
HSPA5
MMP13
ASPN
TUBB
FGFR1
MATN1
P4HB
COL11A1
SOST
APOC1
MFGE8
MSN
POSTN
AFP
VCAN
PCSK9
C3
COL2A1
IGFBP3
ACTB
SPARCL1
SERPINA1
EMILIN1
IGFBP5
IGFBP1
IGLC1
SOD3
HTRA1
IGKC
ALPL
APOA1
ECM2
APP
30
25
20
15
10
5
Petrous - Cortical
Femur - Trabecular
Femur - Growth plate
Rib - Trabecular
Parietal - Cortical Interior
Rib - Cortical
Femur - Cortical
Parietal - Cortical Exterior
Parietal - Trabecular
Parietal - Suture

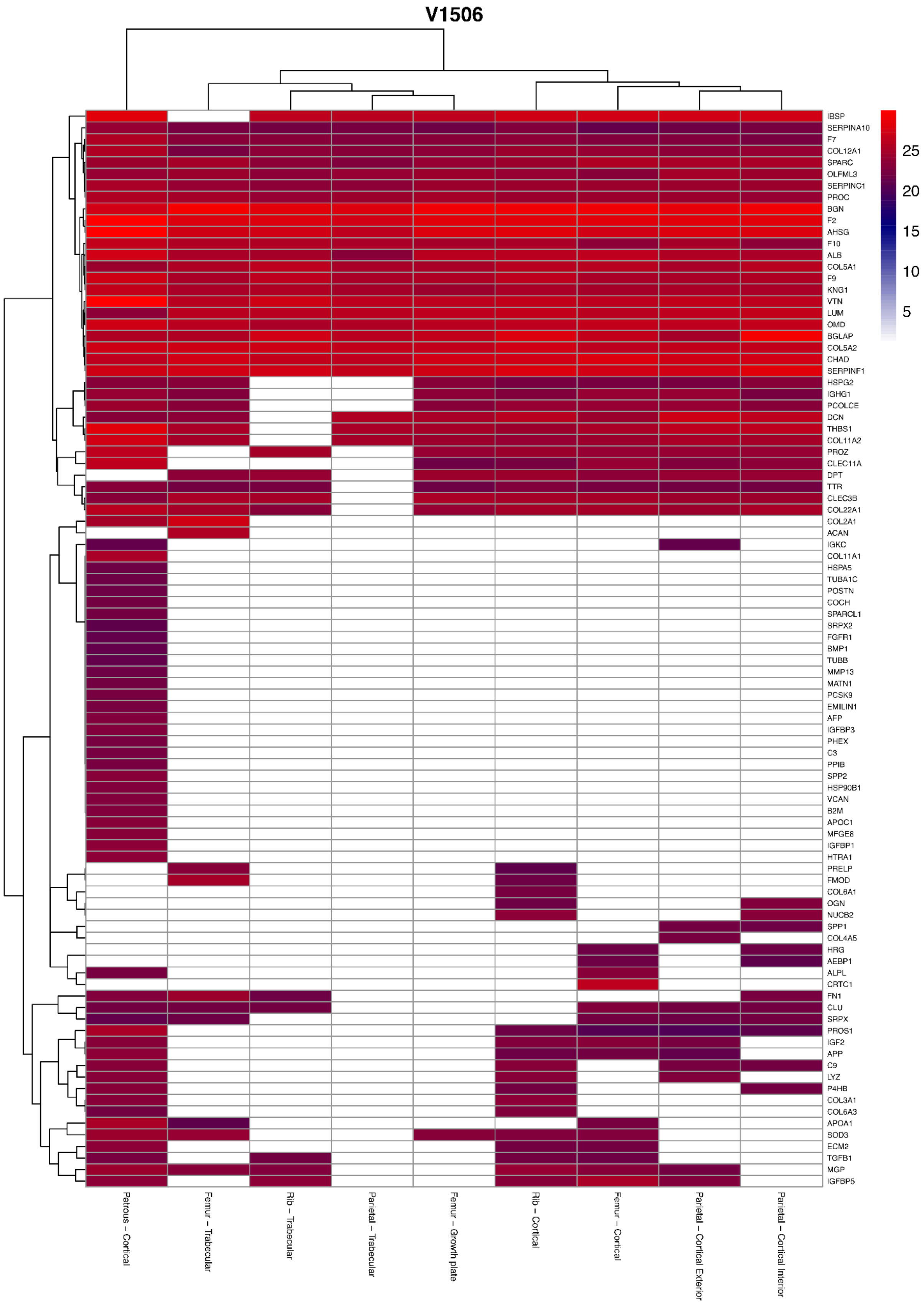
V1506
25
20
15
10
5
IBSP
SERPINA10
F7
COL12A1
SPARC
OLFML3
SERPINC1
PROC
BGN
F2
AHSG
F10
ALB
COL5A1
F9
KNG1
VTN
LUM
OMD
BGLAP
COL5A2
CHAD
SERPINF1
HSPG2
IGHG1
PCOLCE
DCN
THBS1
COL11A2
PROZ
CLEC11A
DPT
TTR
CLEC3B
COL22A1
COL2A1
ACAN
IGKC
COL11A1
HSPA5
TUBA1C
POSTN
COCH
SPARCL1
SRPX2
FGFR1
BMP1
TUBB
MMP13
MATN1
PCSK9
EMILIN1
AFP
IGFBP3
PHEX
C3
PPIB
SPP2
HSP90B1
VCAN
B2M
APOC1
MFGE8
IGFBP1
HTRA1
PRELP
FMOD
COL6A1
OGN
NUCB2
SPP1
COL4A5
HRG
AEBP1
ALPL
CRTC1
FN1
CLU
SRPX
PROS1
IGF2
APP
C9
LYZ
P4HB
COL3A1
COL6A3
APOA1
SOD3
ECM2
TGFB1
MGP
IGFBP5
Petrous – Cortical
Femur – Trabecular
Rib – Trabecular
Parietal – Trabecular
Femur – Growth plate
Rib – Cortical
Femur – Cortical
Parietal – Cortical Exterior
Parietal – Cortical Interior

# V1621

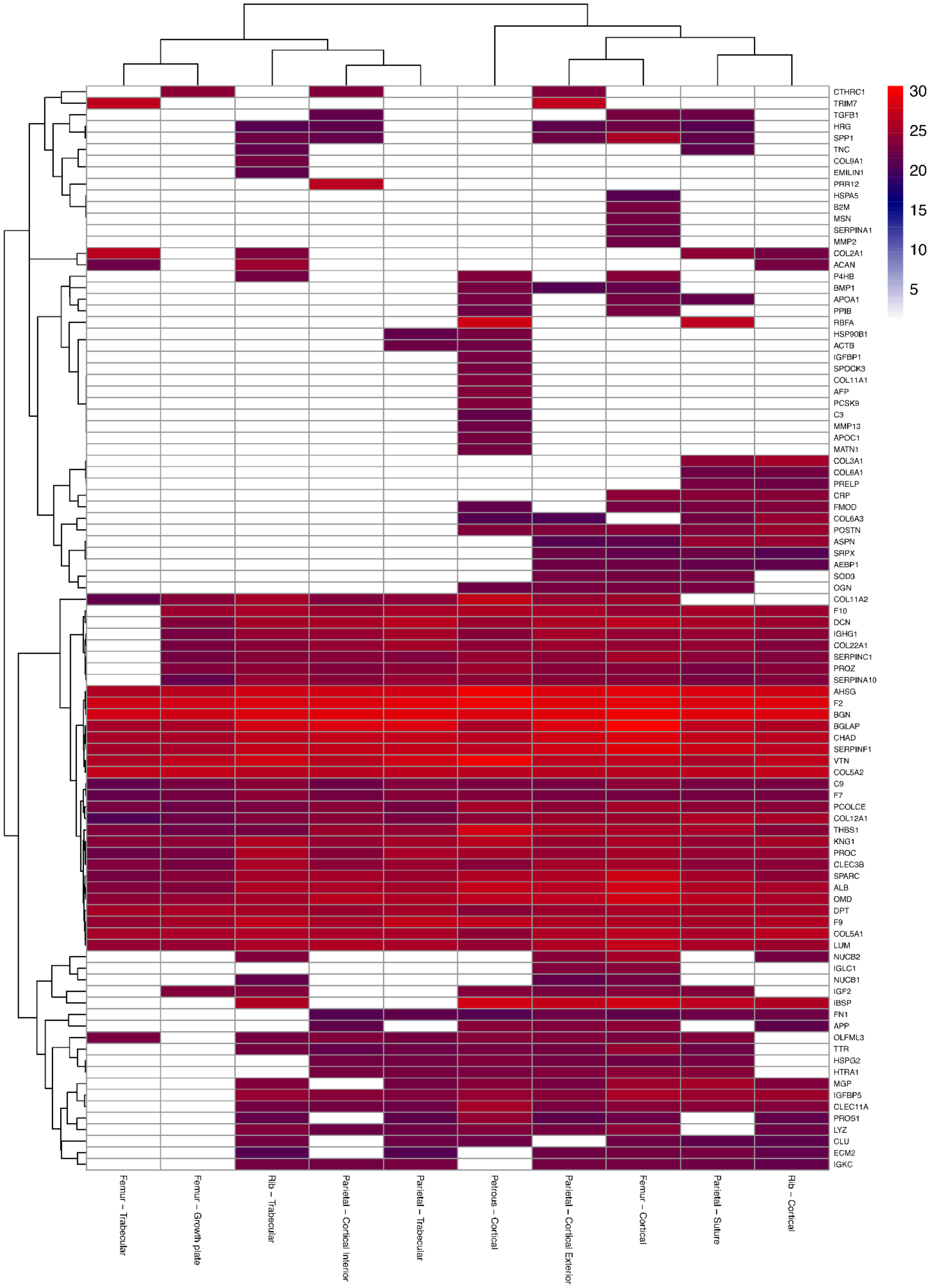

30
25
20
15
10
5
CTHRC1
TRIM7
TGFB1
HRG
SPP1
TNC
COL9A1
EMILIN1
PRR12
HSPA5
B2M
MSN
SERPINA1
MMP2
COL2A1
ACAN
P4HB
BMP1
APOA1
PPIB
RBFA
HSP90B1
ACTB
IGFBP1
SPOCK3
COL11A1
AFP
PCSK9
C3
MMP13
APOC1
MATN1
COL3A1
COL6A1
PRELP
CRP
FMOD
COL6A3
POSTN
ASPN
SRPX
AEBP1
SOD3
OGN
COL11A2
F10
DCN
IGHG1
COL22A1
SERPINC1
PROZ
SERPINA10
AHSG
F2
BGN
BGLAP
CHAD
SERPINF1
VTN
COL5A2
C9
F7
PCOLCE
COL12A1
THBS1
KNG1
PROC
CLEC3B
SPARC
ALB
OMD
DPT
F9
COL5A1
LUM
NUCB2
IGLC1
NUCB1
IGF2
IBSP
FN1
APP
OLFML3
TTR
HSPG2
HTRA1
MGP
IGFBP5
CLEC11A
PROS1
LYZ
CLU
ECM2
IGKC
Femur – Trabecular
Femur – Growth plate
Rib – Trabecular
Parietal – Cortical Interior
Parietal – Trabecular
Petrous – Cortical
Parietal – Cortical Exterior
Femur – Cortical
Parietal – Suture
Rib – Cortical

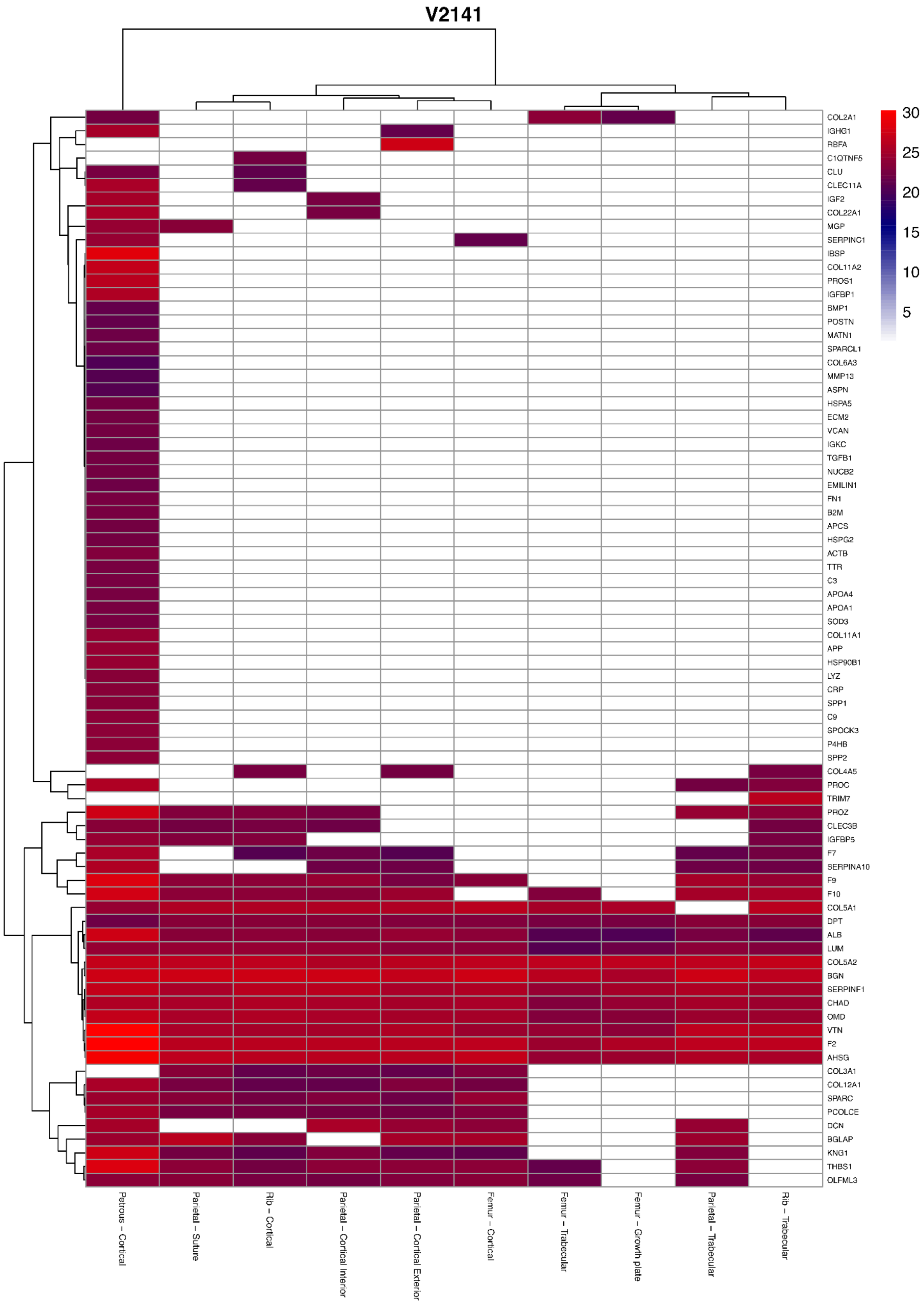

V2141
30
25
20
15
10
5
COL2A1
IGHG1
RBFA
C1QTNF5
CLU
CLEC11A
IGF2
COL22A1
MGP
SERPINC1
IBSP
COL11A2
PROS1
IGFBP1
BMP1
POSTN
MATN1
SPARCL1
COL6A3
MMP13
ASPN
HSPA5
ECM2
VCAN
IGKC
TGFB1
NUCB2
EMILIN1
FN1
B2M
APCS
HSPG2
ACTB
TTR
C3
APOA4
APOA1
SOD3
COL11A1
APP
HSP90B1
LYZ
CRP
SPP1
C9
SPOCK3
P4HB
SPP2
COL4A5
PROC
TRIM7
PROZ
CLEC3B
IGFBP5
F7
SERPINA10
F9
F10
COL5A1
DPT
ALB
LUM
COL5A2
BGN
SERPINF1
CHAD
OMD
VTN
F2
AHSG
COL3A1
COL12A1
SPARC
PCOLCE
DCN
BGLAP
KNG1
THBS1
OLFML3
Petrous - Cortical
Parietal - Suture
Rib - Cortical
Parietal - Cortical Interior
Parietal - Cortical Exterior
Femur - Cortical
Femur - Trabecular
Femur - Growth plate
Parietal - Trabecular
Rib - Trabecular

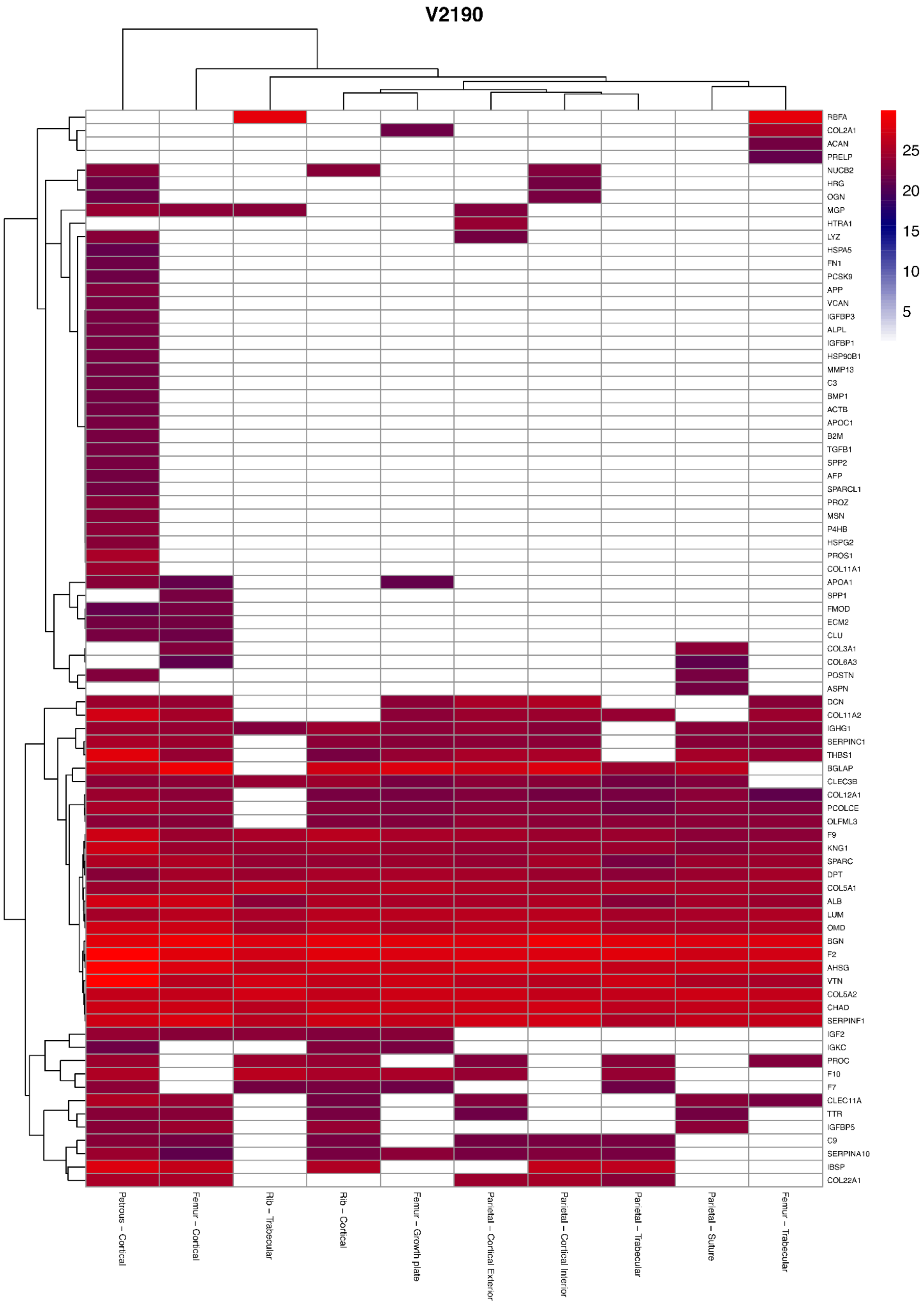
V2190
RBFA
COL2A1
ACAN
PRELP
NUCB2
HRG
OGN
MGP
HTRA1
LYZ
HSPA5
FN1
PCSK9
APP
VCAN
IGFBP3
ALPL
IGFBP1
HSP90B1
MMP13
C3
BMP1
ACTB
APOC1
B2M
TGFB1
SPP2
AFP
SPARCL1
PROZ
MSN
P4HB
HSPG2
PROS1
COL11A1
APOA1
SPP1
FMOD
ECM2
CLU
COL3A1
COL6A3
POSTN
ASPN
DCN
COL11A2
IGHG1
SERPINC1
THBS1
BGLAP
CLEC3B
COL12A1
PCOLCE
OLFML3
F9
KNG1
SPARC
DPT
COL5A1
ALB
LUM
OMD
BGN
F2
AHSG
VTN
COL5A2
CHAD
SERPINF1
IGF2
IGKC
PROC
F10
F7
CLEC11A
TTR
IGFBP5
C9
SERPINA10
IBSP
COL22A1
25
20
15
10
5
Petrous – Cortical
Femur – Cortical
Rib – Trabecular
Rib – Cortical
Femur – Growth plate
Parietal – Cortical Exterior
Parietal – Cortical Interior
Parietal – Trabecular
Parietal – Suture
Femur – Trabecular

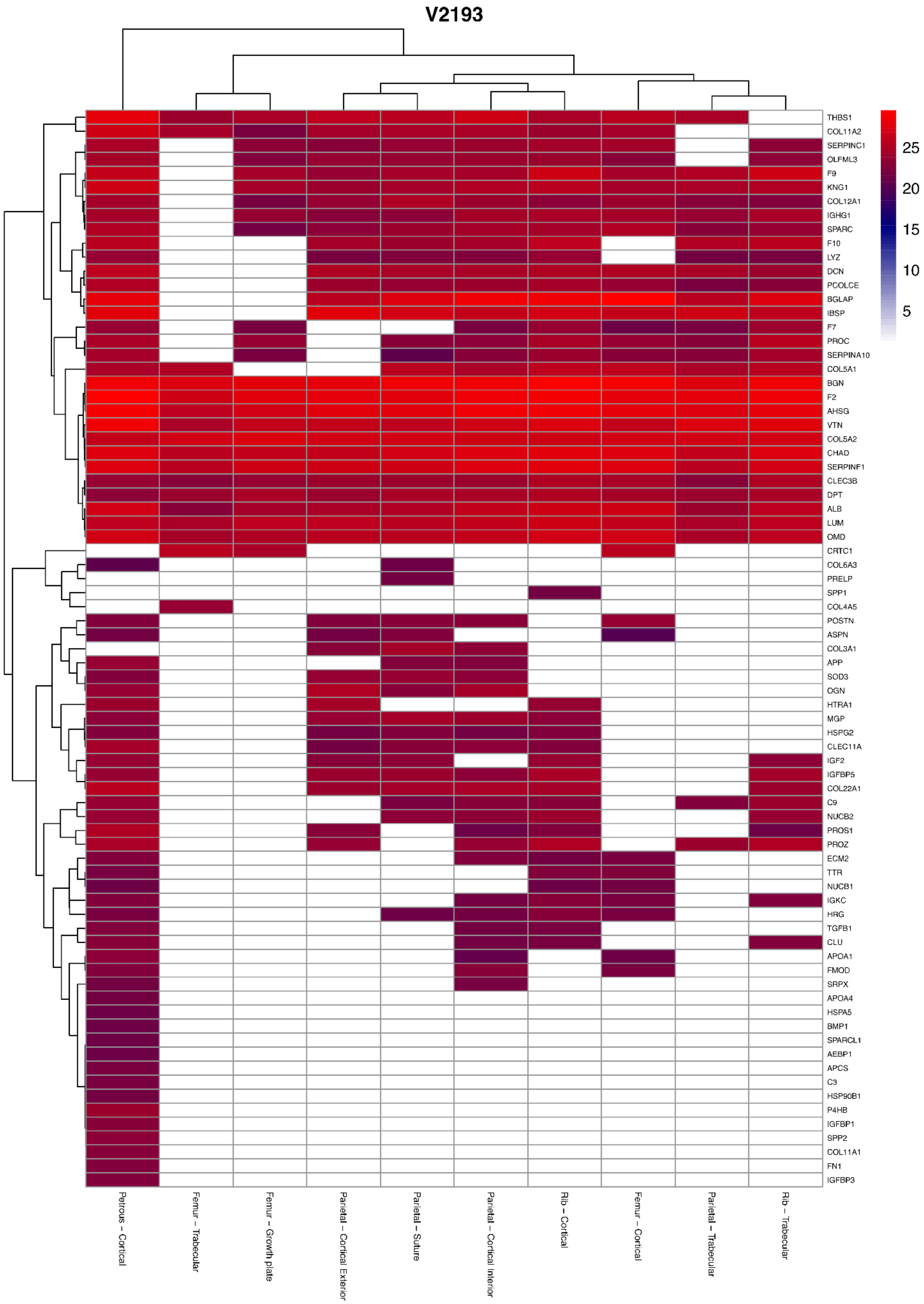

V2193
THBS1
COL11A2
SERPINC1
OLFML3
F9
KNG1
COL12A1
IGHG1
SPARC
F10
LYZ
DCN
PCOLCE
BGLAP
IBSP
F7
PROC
SERPINA10
COL5A1
BGN
F2
AHSG
VTN
COL5A2
CHAD
SERPINF1
CLEC3B
DPT
ALB
LUM
OMD
CRTC1
COL6A3
PRELP
SPP1
COL4A5
POSTN
ASPN
COL3A1
APP
SOD3
OGN
HTRA1
MGP
HSPG2
CLEC11A
IGF2
IGFBP5
COL22A1
C9
NUCB2
PROS1
PROZ
ECM2
TTR
NUCB1
IGKC
HRG
TGFB1
CLU
APOA1
FMOD
SRPX
APOA4
HSPA5
BMP1
SPARCL1
AEBP1
APCS
C3
HSP90B1
P4HB
IGFBP1
SPP2
COL11A1
FN1
IGFBP3
25
20
15
10
5
Petrous – Cortical
Femur – Trabecular
Femur – Growth plate
Parietal – Cortical Exterior
Parietal – Suture
Parietal – Cortical Interior
Rib – Cortical
Femur – Cortical
Parietal – Trabecular
Rib – Trabecular

## V2324

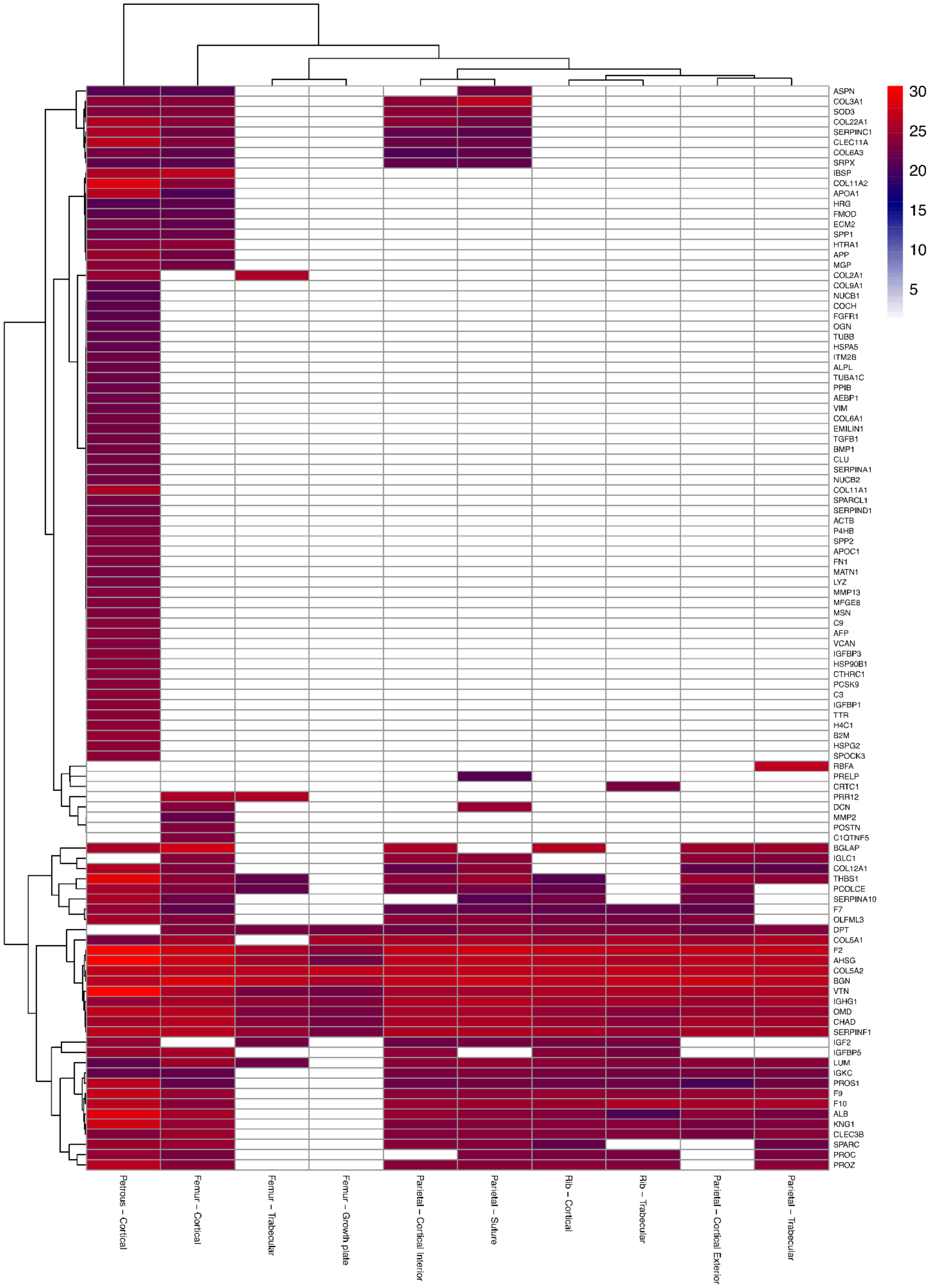

30
25
20
15
10
5
ASPN
COL3A1
SOD3
COL22A1
SERPINC1
CLEC11A
COL6A3
SRPX
IBSP
COL11A2
APOA1
HRG
FMOD
ECM2
SPP1
HTRA1
APP
MGP
COL2A1
COL9A1
NUCB1
COCH
FGFR1
OGN
TUBB
HSPA5
ITM2B
ALPL
TUBA1C
PPIB
AEBP1
VIM
COL6A1
EMILIN1
TGFB1
BMP1
CLU
SERPINA1
NUCB2
COL11A1
SPARCL1
SERPIND1
ACTB
P4HB
SPP2
APOC1
FN1
MATN1
LYZ
MMP13
MFGE8
MSN
C9
AFP
VCAN
IGFBP3
HSP90B1
CTHRC1
PCSK9
C3
IGFBP1
TTR
H4C1
B2M
HSPG2
SPOCK3
RBFA
PRELP
CRTC1
PRR12
DCN
MMP2
POSTN
C1QTNF5
BGLAP
IGLC1
COL12A1
THBS1
PCOLCE
SERPINA10
F7
OLFML3
DPT
COL5A1
F2
AHSG
COL5A2
BGN
VTN
IGHG1
OMD
CHAD
SERPINF1
IGF2
IGFBP5
LUM
IGKC
PROS1
F9
F10
ALB
KNG1
CLEC3B
SPARC
PROC
PROZ
Petrous – Cortical
Femur – Cortical
Femur – Trabecular
Femur – Growth plate
Parietal – Cortical Interior
Parietal – Suture
Rib – Cortical
Rib – Trabecular
Parietal – Cortical Exterior
Parietal – Trabecular

# V2351

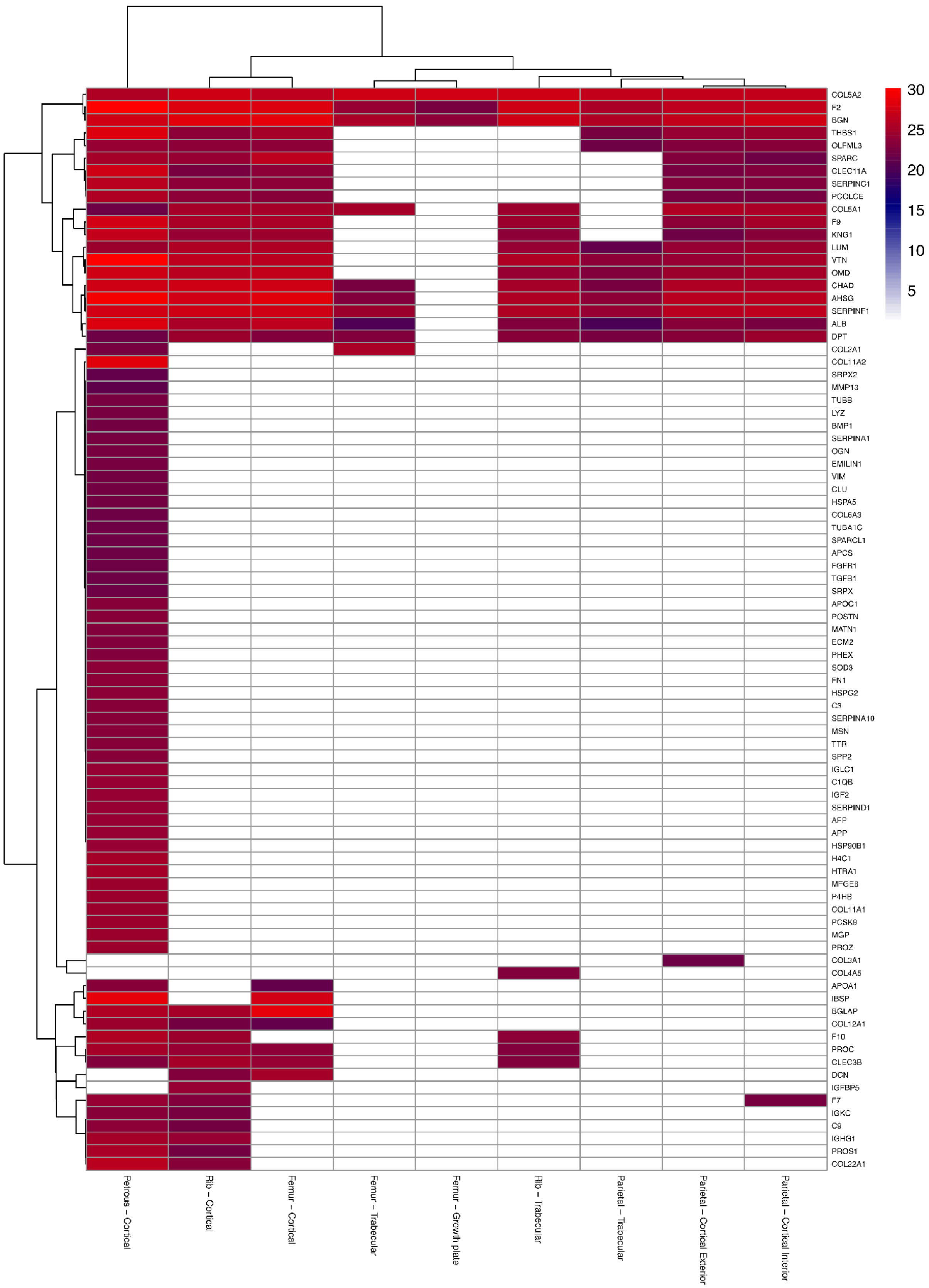

30
25
20
15
10
5
COL5A2
F2
BGN
THBS1
OLFML3
SPARC
CLEC11A
SERPINC1
PCOLCE
COL5A1
F9
KNG1
LUM
VTN
OMD
CHAD
AHSG
SERPINF1
ALB
DPT
COL2A1
COL11A2
SRPX2
MMP13
TUBB
LYZ
BMP1
SERPINA1
OGN
EMILIN1
VIM
CLU
HSPA5
COL6A3
TUBA1C
SPARCL1
APCS
FGFR1
TGFB1
SRPX
APOC1
POSTN
MATN1
ECM2
PHEX
SOD3
FN1
HSPG2
C3
SERPINA10
MSN
TTR
SPP2
IGLC1
C1QB
IGF2
SERPIND1
AFP
APP
HSP90B1
H4C1
HTRA1
MFGE8
P4HB
COL11A1
PCSK9
MGP
PROZ
COL3A1
COL4A5
APOA1
IBSP
BGLAP
COL12A1
F10
PROC
CLEC3B
DCN
IGFBP5
F7
IGKC
C9
IGHG1
PROS1
COL22A1
Petrous – Cortical
Rib – Cortical
Femur – Cortical
Femur – Trabecular
Femur – Growth plate
Rib – Trabecular
Parietal – Trabecular
Parietal – Cortical Exterior
Parietal – Cortical Interior

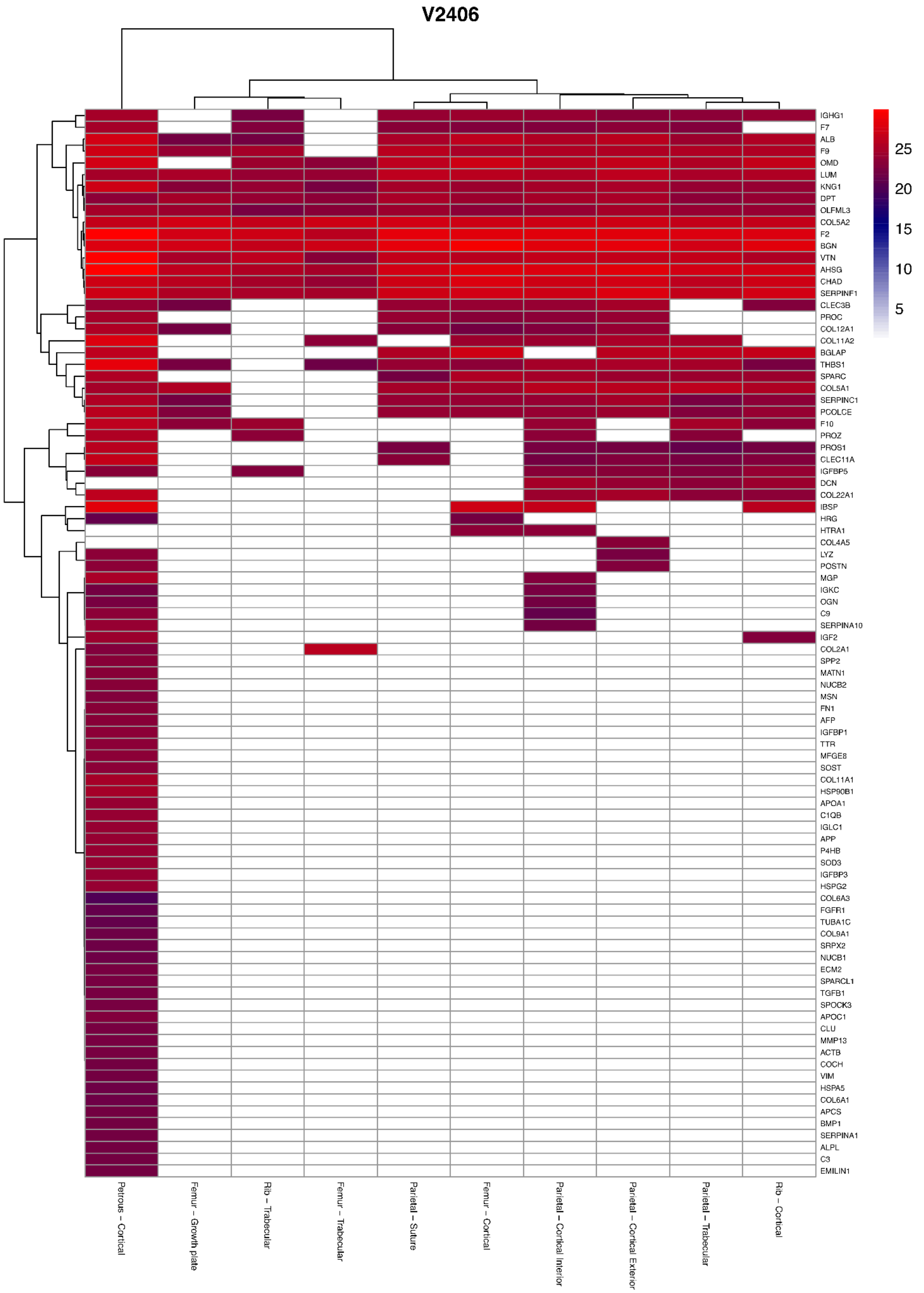
V2406
IGHG1
F7
ALB
F9
OMD
LUM
KNG1
DPT
OLFML3
COL5A2
F2
BGN
VTN
AHSG
CHAD
SERPINF1
CLEC3B
PROC
COL12A1
COL11A2
BGLAP
THBS1
SPARC
COL5A1
SERPINC1
PCOLCE
F10
PROZ
PROS1
CLEC11A
IGFBP5
DCN
COL22A1
IBSP
HRG
HTRA1
COL4A5
LYZ
POSTN
MGP
IGKC
OGN
C9
SERPINA10
IGF2
COL2A1
SPP2
MATN1
NUCB2
MSN
FN1
AFP
IGFBP1
TTR
MFGE8
SOST
COL11A1
HSP90B1
APOA1
C1QB
IGLC1
APP
P4HB
SOD3
IGFBP3
HSPG2
COL6A3
FGFR1
TUBA1C
COL9A1
SRPX2
NUCB1
ECM2
SPARCL1
TGFB1
SPOCK3
APOC1
CLU
MMP13
ACTB
COCH
VIM
HSPA5
COL6A1
APCS
BMP1
SERPINA1
ALPL
C3
EMILIN1
25
20
15
10
5
Petrous – Cortical
Femur – Growth plate
Rib – Trabecular
Femur – Trabecular
Parietal – Suture
Femur – Cortical
Parietal – Cortical Interior
Parietal – Cortical Exterior
Parietal – Trabecular
Rib – Cortical

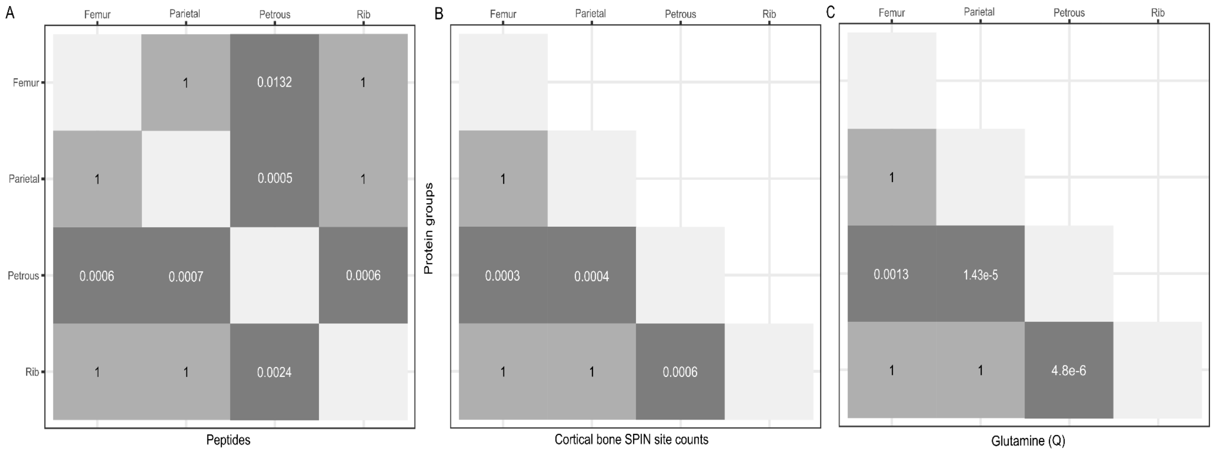


SI Figure 4: Post-hoc tests. A) Comparisons of number of identified protein groups and number of peptides for cortical bone samples. Dunn's test. B) SPIN site counts in cortical bone. Tukey's HSD. C) Glutamine (Q) deamidation of cortical bone samples. Tukey's HSD.

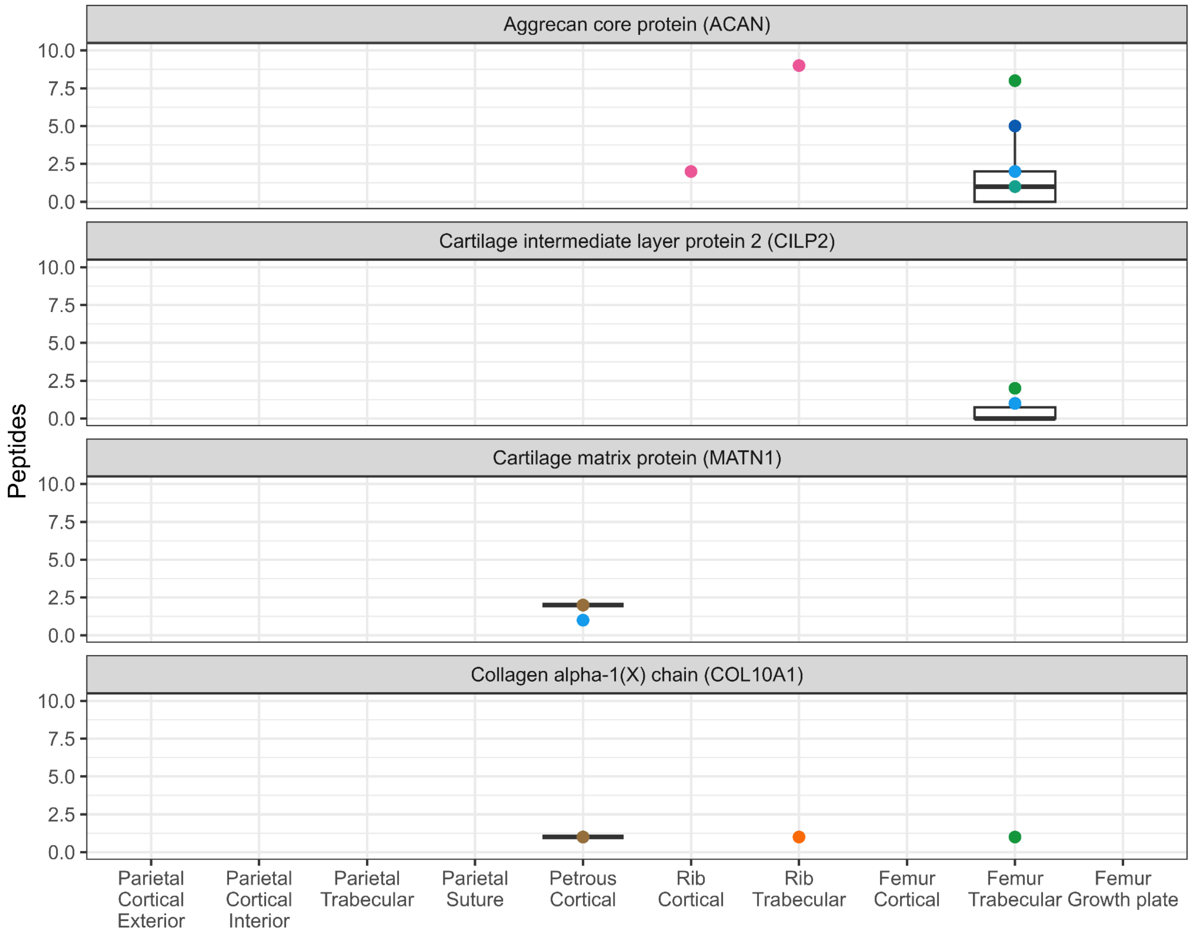


SI Figure 5: Abundance of proteins unique to chondrocytes, expressed during endochondral ossification. Colour of points indicating individuals, and are the same as in SI Figure 1. An absence of points implies that no peptides were identified in a particular extraction.
